# Form IF Rubiscos include highly active, specific, and small subunit-independent enzymes

**DOI:** 10.64898/2026.03.18.712719

**Authors:** F. Otto, H. Westedt, K. P. Franzeck, J. Zarzycki, A. M. Küffner, L. Schulz, S. Prinz, N. Paczia, P. Claus, G. K. A. Hochberg, T. J. Erb

## Abstract

Plant-type (Form I) Ribulose-1,5-bisphosphate carboxylase/oxygenase (Rubisco) suffers from inherent catalytic trade-offs and a strong dependency on other proteins—including an essential small subunit (SSU) and auxiliary chaperones—for assembly, constraining the enzyme’s evolutionary and engineering potential. Here, we investigated representatives from the newly discovered clade Form IF. These enzymes do not require specific chaperones to form functional complexes, exhibit high CO_2_-specificities (S_C/O_ ∼50) while maintaining high turnover rates (up to *k*_cat_ ∼11 s^−1^). Remarkably, two Form IF representatives (IF-1/IF-2) lost the dependency on the SSU and assemble into homo-octameric complexes without their cognate SSUs. While the SSU is not necessary for catalysis, its addition improves both activity and specificity in IF-1/IF-2. Our results show that complexity is actually not required to achieve highly active, specific and functional Rubisco variants—and that this complexity can even be reverted—which challenges our current thinking on the evolution and catalytic mechanism of Rubisco.

## Introduction

The enzyme Ribulose-1,5-bisphosphate Carboxylase/Oxygenase (Rubisco) plays a pivotal role in the biological carbon cycle, where it catalyzes the assimilation of more than 400 Gt CO₂ annually within the Calvin-Benson-Bassham (CBB) cycle^1–3^. Rubisco is the only carboxylase that suffers from an inherent oxygenase side activity, which can be as high as 25 % at ambient air, resulting in the energetically costly process of photorespiration, representing a limiting factor in photosynthesis^4–6^ and additionally displays a relatively low catalytic turnover rate (*k*_cat_ 1 – 10 s^−1^)^7^. To overcome these catalytic limitations of Rubisco, many studies focused on engineering the enzyme towards increased activity and/or specificity^1,8–15^. Critically, many of these efforts show that when attempting to increase Rubiscos catalytic rate, the enzyme becomes less specific and *vice versa*^1,16,17^. This apparent trade-off between activity and specificity of Rubisco has become an intense research subject in mechanistic enzymology and evolutionary biochemistry^8–11,18–22^.

The Rubisco enzyme family has a rich evolutionary history that dates back several billions of years. Rubisco homologs are found across diverse phylogenetic lineages and exist in four different subtypes (Form I-IV) that assemble in different oligomeric states with distinct biochemical properties^23^. Rubiscos Form II, III and IV are primarily found in bacteria and archaea. They consist of large subunits (LSUs) that form the catalytically active core as homo-dimers (L_2_), and higher order assemblies (L_4_ – L_10_). In contrast, Form I Rubiscos, which are the dominant Rubisco form on Earth, and found in plants, mosses and (cyano)bacteria are more complex^24^. These enzymes exist as hetero-hexadecamers composed of eight LSUs that form a catalytic core and eight small subunits (SSUs), which cap off the LSUs, resulting in an L_8_S_8_ assembly.

Notably, the acquisition of the SSU during evolution seems to have benefitted several catalytic parameters of the core enzyme, including CO_2_ specificity. However, it also introduced a critical dependency of the LSU on the SSU^18,25^. This dependency emerged relatively early in the evolution of Form I Rubiscos, prior to their splitting into five different clades that encompass clades IA/B (“green type” Form I), clades IC/D (“red type” Form I) and the recently discovered clade IF (Figure 1 A)^18,25,26^. So far, all Rubiscos from within these clades require co-expression with their SSU to ensure proper folding, stability, and activity^27,28^.

**Figure 1.**
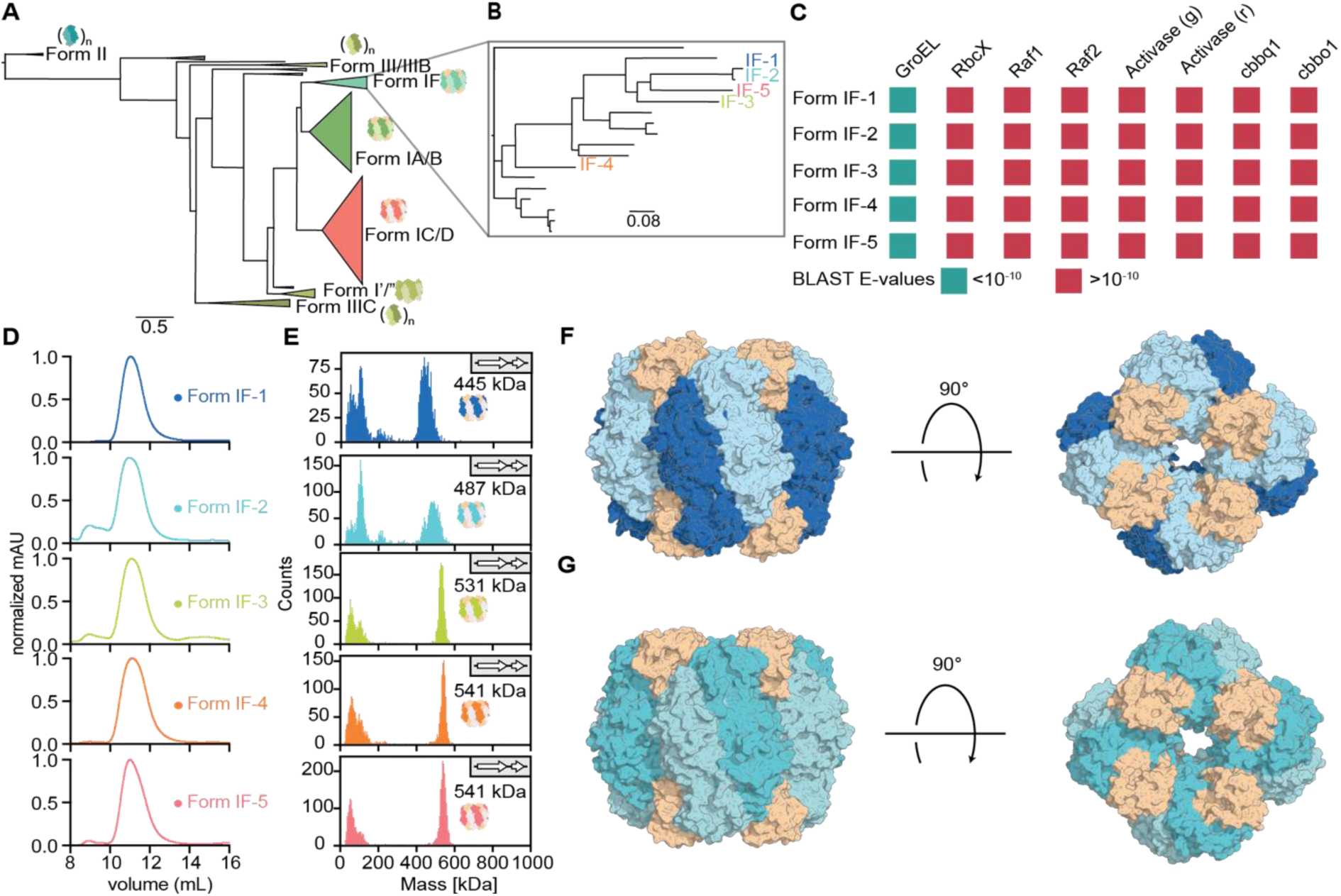
Form IF Rubiscos form a separate clade in the phylogeny and do not require specific chaperones for assembly into functional L_x_S_x_ complexes. (A) Reduced phylogenetic tree of Rubisco with the Form IF Rubiscos clade (teal), branching from Form IA/B Rubiscos highlighted in green. (B) Phylogenetic tree of the Form IF clade. Proteins of interest are denoted on the tree. Form IF-1 and IF-2 belong to the *Gaiellales* bacterium clade, Form IF-3 can be found in a *Chloroflexi* bacterium, Form IF-4 in an *Anaerolineales* bacterium and Form IF-5 in a *Limnochordaceae* bacterium. (C) Metagenomes of the hosts of the proteins of interest were searched for known Rubisco chaperones and activases. Dark green denotes presence of a gene encoding a homolog (with a BLAST E-value of <10^−10^, red fields denote the lack of a gene encoding a homolog). None of the hosts harbor enzymes like known Rubisco chaperones. (D) Size-exclusion chromatography revealed the hexadecameric complex, as indicated by the calculated L_8_S_8_ retention volume. (E) Mass Photometry showing L_8_S_8_ formation for Form IF-3 (green) Form IF-4 (orange), Form IF-5 (coral) and mixed results of L_8_/L_8_S_8_ formation for Form IF-1 (dark blue) and Form IF-2 (turquoise). (F) Cryo-EM structure of IF-1. The LSU is shown in blue and light blue, the SSU in beige. IF-1 PDB: 28VR. (G) Crystal structure of IF-2. LSU is shown turquoise and light turquoise the SSU in ocher. IF-2 PDB: 28VS.

Thus, while the addition of the SSU allowed Form I Rubiscos to explore a different sequence space to fine-tune their catalytic properties (e.g., improved specificity)^12,28^, it came at further costs^25,29^. In particular, the functional interdependence between LSU and SSU imposed additional requirements for protein folding and assembly. Well-studied Form IA/B Rubiscos rely not only on the SSU but also on a suite of chaperones, including GroEL/GroES (Cpn60/Cpn10 in plants), RbcX, Raf1, and Raf2 for maturation and long-term operation^12,30–34^. These data have led to the current model of the “Rubiscosome”, in which modern-day, high-performance Form I Rubiscos have become fully dependent on the SSU and multiple associated proteins^17,35,36^.

Here, we describe several homologs from the IF clade that challenge the idea that these interactions are strictly essential for Form I Rubiscos. We characterized different Form IF Rubiscos to identify homologs that are chaperone-independent and fully functional in the absence of their SSU. These enzymes form comparatively “simple” L_8_ complexes that exhibit high specificity (S_C/O_ ∼40) and specific activity (*k*_cat_ 4.3 s^−1^), resembling the traits of their more complex plant counterparts. Apparently, these IF Rubiscos have lost the functional dependency on their SSU^37^, which represents a unique event in Rubisco evolution and opens new possibilities for creating or transplanting high-performing Rubiscos independent of additional protein components.

## Results

### Form IF Rubiscos include highly specific and active enzyme variants

Recently, a new subclade of Form I Rubisco was discovered from metagenomics studies^18,26^. These Form IF Rubiscos are phylogenetically positioned between Form IA/B and Form IC/D clades and consist of around 20 known representatives (Figure 1 A/B). According to our analysis (Figure 1 C), all host genomes that encode the enzymes of our interest encode a LSU and a SSU but notably lack clear homologs of Rubisco chaperones and/or activases, suggesting that IF enzymes might not require specific assistance for folding. To investigate the properties of this previously uncharacterized clade of enzymes, we focused on five representatives, designated Form IF-1 through IF-5 (Figure 1B).

All five homologs were co-expressed as LSU-SSU constructs (indicated by subscript _LS_) and resulted in soluble, enzymatically active proteins. Size-exclusion chromatography (SEC) and mass photometry indicated that all enzymes adopted higher order complexes (L_X_S_X_), consistent with Form I Rubisco assemblies (Figure 1 D/E). Interestingly, while Form IF-1_LS_ and IF-2_LS_ showed enzymatic activity, mass photometry indicated that they did not form fully occupied L_8_S_8_ complexes (Figure 1 E), suggesting that complete SSU occupancy may not be strictly required for catalytic function in these variants. Next, we studied the kinetic parameters of the five IF representatives (Table 1). All enzymes catalyzed ribulose-1,5-bisphosphate carboxylation at turnover rates (*k*_cat_) between 0.7 s^−1^ (IF-4_LS_ & IF-5_LS_) and up to 6.9 s^−1^ (IF-1_LS_ & IF-2_LS_), which resembles the range of known plant homologs. The CO_2_ *K*_m_ values of these five enzymes varied from 40 µM for (IF-1_LS_) to 237 µM CO_2_ (IF-4_LS_). The specificity for CO_2_ over O_2_ (S_C/O_) ranged from 23 to 53 (Table 1), which is comparable to values reported for cyanobacterial Rubiscos^20^. Altogether, these data showed that (most) clade IF enzymes are highly specific and that some representatives (IF-1_LS_ & IF-2_LS_) possess additional high activities (k_cat_ 6.9 s^−1^), positioning them among the best-performing plant variants characterized to date (Figure 2 J)^14,20,38^.

**Figure 2.**
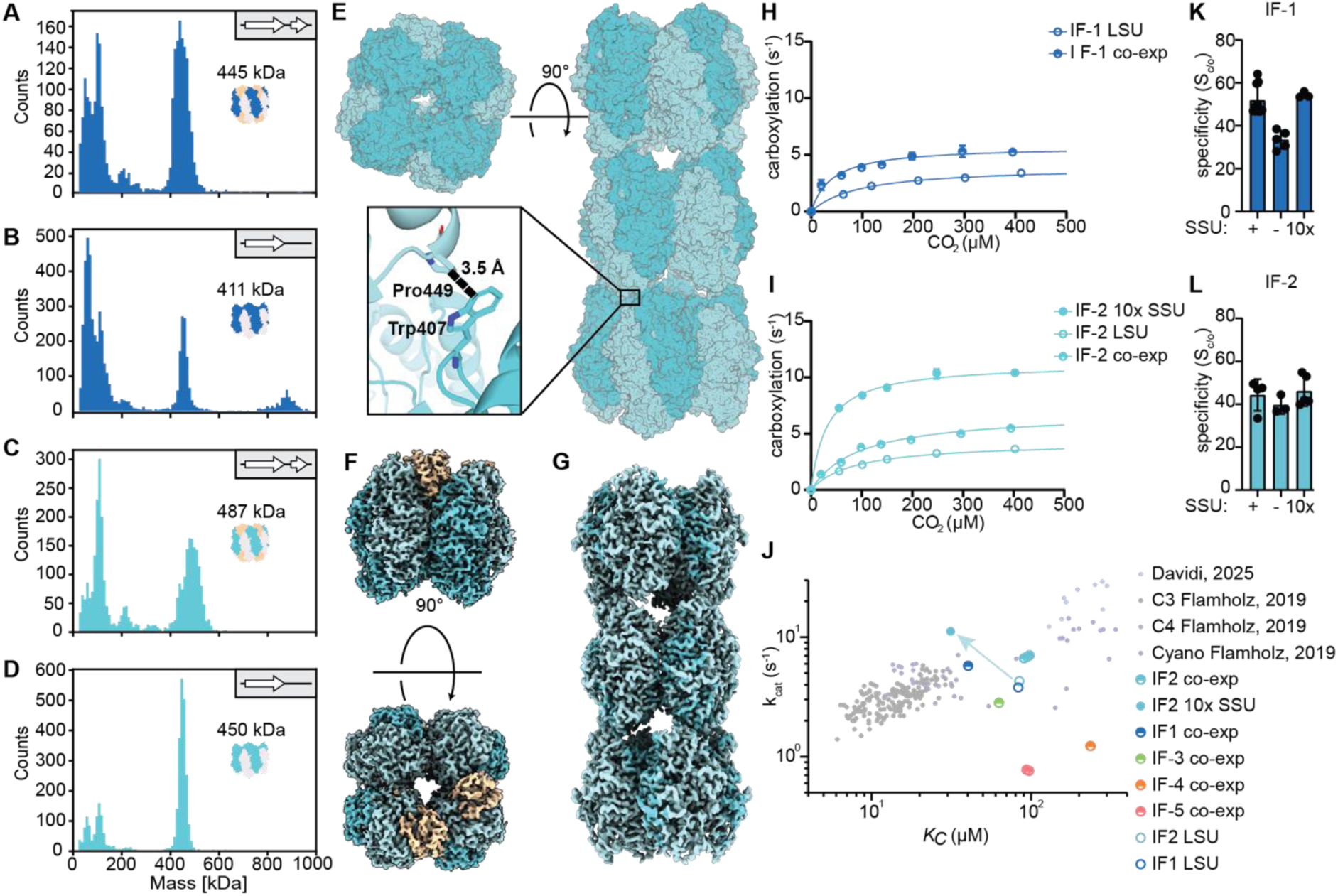
IF-1 and IF-2 function independent of the SSUs but show improved kinetics upon SSU addition. (**A**) Mass photometric analysis of co-expressed IF-1 Rubisco. The peak at 445 kDa does not correspond to the expected weight of a L_8_S_8_ complex but rather a complex that is not fully occupied with the SSU. (**B**) Mass photometric analysis of LSU-only IF-1_L_ Rubisco with the main peak displaying the expected weight for a L_8_ LSU-only construct. 21 % of the sample can be observed as a L_8_ complex at a concentration of 25 nM. (**C**) Mass photometric analysis of co-expressed IF-2 Rubisco. The peak at 487 kDa does not correspond to the expected weight of L_8_S_8_ complex but rather a complex that is not fully occupied with the SSU. (**D**) Mass photometric analysis of IF-2 LSU-only Rubisco with the main peak displaying the expected weight for a L_8_ LSU-only construct of 450 kDa. 64 % of the sample can be observed as a L_8_ complex at a concentration of 25 nM. (**E**) Cryo-EM analysis of the IF-2 LSU-only complex forming fibrils by stacking *via* the SSU binding positions. The binding interaction between Pro449 and Trp407 of different LSUs that likely leads to the formation fibrils is shown in a close-up view. (**F**) Cryo-EM density map of IF-2 Rubisco shows the lacking SSU density. (**G**) Cryo-EM electron density map of the IF-2_L_ complex. Shown here is a triple stack as a part of a twisted fibril. PDB: 28VT. (**H**) Turnover rate (*k*_cat_) of IF-1 as LSU-only construct and in standard co-expression assembly. (**I**) Turnover rate (*k*_cat_) of IF-2_L_, in standard co-expression assembly and with 10-fold excess of the SSU. Notably, the 10-fold molar excess of the SSU leads to drastic increase in carboxylation rate for IF-2. (**J**) Influence of SSU stoichiometries onto turnover rate and apparent CO_2_ specificities (*K*_m_ CO_2_) of IF-1 and IF-2. IF-1 to IF-5 as co-expressed enzymes and previously characterized Rubiscos from literature are displayed as comparisons^8,16^. The arrow in turquoise highlights the difference between IF-2 LSU-only and IF-2 with 10-fold excess of the SSU. (**K**) Specificities (S_C/O_) of IF-1 without SSU and at 10-fold excess of the SSU. (**L**) Specificities (S_C/O_) of IF-2 without SSU and at 10-fold excess of the SSU.

**Table 1.** Kinetic parameters of Form IF Rubiscos and selected other Rubiscos. *k*_cat_ is the maximal rate of carboxylation under saturating substrate concentrations at 25 °C. *K*_m_(CO2) is the Michaelis constants for CO_2_. S_C/O_ = [*k*_cat_C/*K*_m_(CO2)]/[*k*_cat_O/*K*_m_(O_2_)]. Values are means ± 95% confidence intervals, ND not determined. *A. thaliana, Arabidopsis thaliana; P. breve Promineofilum breve*.

| Rubisco | $k_{cat}$ (s<sup>-1</sup>) | $K_m$ (CO<sub>2</sub>) (μM) | Specificity (S<sub>C/O</sub>) | $k_{cat}/K_m$ (CO<sub>2</sub>) (M<sup>-1</sup>s<sup>-1</sup>) |
| --- | --- | --- | --- | --- |
| IF-1<sub>LS</sub> | 5.8 ± 0.5 | 40.4 ± 6.2 | 51.9 ± 6.7 | 1.42 × 10<sup>5</sup> |
| IF-2<sub>LS</sub> | 6.9 ± 0.2 | 94.3 ± 3.0 | 52.7 ± 6.8 | 7.28 × 10<sup>4</sup> |
| IF-3<sub>LS</sub> | 2.8 ± ND | 63.1 ± ND | 41.2 ± 1.3 | 4.48 × 10<sup>4</sup> |
| IF-4<sub>LS</sub> | 1.2 ± ND | 237 ± ND | 24.7 ± 4.6 | 5.20 × 10<sup>3</sup> |
| IF-5<sub>LS</sub> | 0.8 ± ND | 95.1 ± ND | 37.0 ± 0.7 | 8.06 × 10<sup>3</sup> |
| IF-1<sub>L</sub> (LSU only) | 3.8 ± 0.3 | 83.3 ± 14.7 | 33.5 ± 2.4 | 4.55 × 10<sup>4</sup> |
| IF-2<sub>L</sub> (LSU only) | 4.3 ± 0.2 | 84.5 ± 7.42 | 39.7 ± 4.2 | 5.04 × 10<sup>4</sup> |
| IF-1<sub>L</sub> + 10 × SSU | ND | ND | 54.3 ± 1.4 | ND |
| IF-2<sub>L</sub> + 10 × SSU | 11.2 ± 0.4 | 31.4 ± 2.6 | 46.3 ± 7.1 | 3.58 × 10<sup>5</sup> |
| *A. thaliana* (form IB) | 3.3 ± 0.1 | 11.8 ± 0.5 | 75.1 ± 0.1 | 2.81 × 10<sup>5</sup> |
| *P. breve* (form I') | 2.2 ± 0.0 | 22.2 ± 9.7 | 36.1 ± 0.9 | 1.00 × 10<sup>5</sup> |

### IF-1 and IF-2 are not dependent on SSU for Rubisco activity

Because of their high activity and specificity within the IF clade, we aimed to further investigate the IF-1 and IF-2 LSU-SSU enzyme complexes in more detail. Closer inspection of our mass photometry data indicated incomplete SSU occupancy of the purified IF-1_LS_ and IF-2_LS_ complex, respectively and suggested that both enzymes predominantly formed an L_8_S_3_ oligomer (Figure 1 E, Figure 2 A). These observations were independently confirmed by cryo-EM of the purified LSU-SSU construct of IF-1 (Figure 1 F, Figure S 2) and IF-2 Rubisco. For IF-2 cryo-EM confirmed assembly of the LSU into an octameric core, but revealed reduced electron density at the SSU-LSU interface, suggesting incomplete SSU incorporation (Figure 2 G, Figure S 4), while the crystal structure at higher protein concentrations showed a hetero-hexadecameric structure (Figure 1 G). Altogether, this data led us to hypothesize that IF-1 and IF-2 Rubisco might not be strictly dependent on their SSU.

Indeed, when expressing the LSUs of IF-1 and IF-2 in the absence of their cognates SSUs (“LSU-only”, indicated by subscript _L_), we obtained soluble protein that formed stable L_8_ complexes (Figure S 1 B/C). Surprisingly, Form IF-1L and IF-2 L assemblies retained catalytic activity, despite absence of their cognate SSUs (Figure 2). Mass photometry analysis showed that the majority of IF-1L and IF-2L particles assembled into L8 complexes (Figure 2 B/D), which is in line with enzymes from the Form I’ clade that do not possess a SSU 33. For IF-2L, the formation of homo-octameric complexes was additionally confirmed by cryo-EM (Figure *2* E/G, Figure S 3), which additionally showed assembly of some L8 complexes of IF-2L into higher ordered structures, i.e. L8-fibrils.

More surprisingly, when assessing the kinetic parameters of IF-1_L_ and IF-2_L_, we observed that the LSU-only complexes maintained high carboxylation rates (*k*_cat_ ∼ 4 s^−1^, Figure 2 H/I) without major changes in specificity (S_C/O_ ∼40; Figure 2 K/L), showing that Form IF enzymes can be active and specific in the absence of their cognate SSUs.

Having characterized the basic properties of the LSU-only complex, we next investigated the impact of forced full SSU occupancy on the catalytic properties of Form IF-2 Rubisco. To that end, we reconstituted the L_8_ core with a tenfold molar excess of its native SSU. This supplementation resulted in a three-fold more active complex (Figure 2 I) compared to IF-2_L_ (*k*_cat_ 11.2 s^−1^), and also increased specificity of the enzyme to an S_C/O_ of 46 (54 for IF-1), highlighting a strong contribution of the SSU to key catalytic properties of the enzyme (Figure 2 K/L, Table 1).

### Single-point mutations introduce SSU-dependency into IF-1 & IF-2

In contrast to IF-1 and IF-2, all other representatives of clade IF (IF-3 to IF-5) could not be produced as soluble, active proteins without their cognate SSU, indicating that IF-1 and IF-2 represent unique cases within the Form I clade that regained the ability to function in the absence of the SSU. To study key mutations along this potential evolutionary trajectory, we focused on comparing IF-1 and IF-2, which are phylogenetically closely related, to their closest characterized representatives IF-5, which is dependent on the SSU for solubility.

Sequence analysis in combination with inspection of predicted structures identified several amino acids at the LSU/SSU interface that differ between IF-1/2 and IF-5, potentially responsible for SSU-dependence (Figure 3 A). To test the effect of those residues on solubility, we tried to induce SSU-dependency in IF-1 by partial substitution of interface residues into their IF-5 counterparts. We chose to probe positions V71T, Q152V and R179K (IF-1 numbering), which are all located at the SSU/LSU interface. Furthermore, we also probed position K425 due to its surface-exposed position on the LSU/SSU interface. All of the tested IF-1 mutants were insoluble in absence of the SSU but not when the SSU was co-expressed (Figure 3 C), reconfirming that the SSU is capable of buffering deleterious substitutions^25^. With dependency-conferring residues identified, we next sought to make IF-5 SSU-independent by substitution of those residues into their IF-1 equivalent. However, none of the IF-5 single-mutants became soluble in absence of their SSU. In line with previous studies on ancestral Rubiscos, one single mutation can cause the SSU to become essential while reversion to an independent state is hardly possible once a SSU-dependency was gained^25^. Even though more work is needed to elucidate IF-1 and IF-2’s evolutionary trajectories, they, contrary to all other so far characterized high-specificity Form I Rubiscos, seem to only be entrenched for catalysis, but not for solubility.

**Figure 3.**
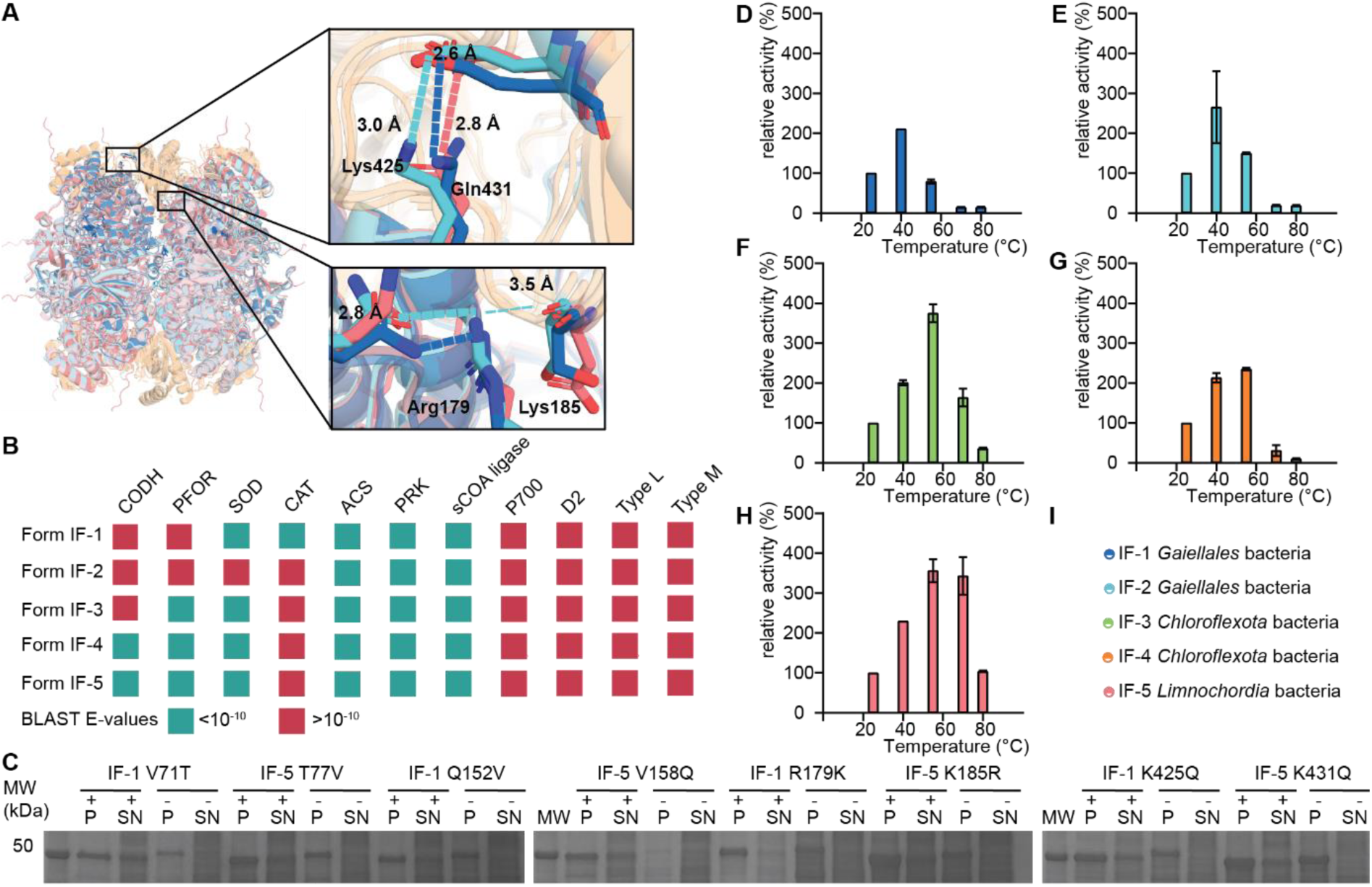
Evolutionary and physiological trends within clade IF Rubiscos. (**A**) Amino acid differences at the SSU/LSU interface between IF-1& IF-2 (SSU-independent for solubility) and closely related IF-5 (SSU-dependent for solubility). The residues in IF-1 R179 and K185 are located on the right side of the LSU/SSU interface. The residues K425 and Q431 are located on the left side of the cleft. IF-1 PDB: 28VR, IF-2 PDB: 28VS. IF-5 is depicted as an alphafold2 prediction^39^. (**B**) Metagenomic analysis of host organisms of Form IF Rubisco for photosynthetic and (an)aerobic lifestyle. Host organisms of IF-1 to IF-5 were searched for specific enzymes that give indications about aerobic or anaerobic lifestyle of those organisms. CODH = carbon monoxide dehydrogenase, PFOR = pyruvate:ferredoxin oxidoreductase, SOD = superoxide dismutase, CAT = catalase, TALDO = transaldolase, PRK = phosphoribulokinase. P700 was used as key enzyme to check of the presence of photosystem I. The D2 protein of photosystem II was used as its respective marker. Green colored fields denote presence of the respective protein within the hosts’ genome; red fields denote lack of a gene encoding the respective protein. The E-value of >10^−10^ was used as a cutoff to discard a gene as encoding a clear homolog. (**C**) Solubility analysis of SSU/LSU interface variants of IF-1 & IF-5. Introduction of residues from the SSU-dependent IF-5 results in failure to assemble of the IF-1 LSU-only complex. Introduction of residues from the SSU-independent IF-1 does not result assembly of an IF-5 LSU-only complex. MW: molecular weight in kDa, P: pellet, SN: supernatant, +/- indicating the presence of the respective SSU. (**D**) – (**H**) Temperature dependent relative activities to their activity at 25°C (see Table 1 for the activities at 25°C) for Form IF-1 to IF-5. (**I**) Listed are the organism from Form IF-1 to IF-5 Rubiscos. *Chloroflexa* is considered as an aerobic thermophile, *Limnochordia* is considered as thermophilic bacteria.

### Metagenomics data to elucidate Rubiscos function and prevalence

Having shown that the SSU affects both, activity, and specificity of the core enzyme, is in line with earlier data^18^ and raised the question, whether enhanced activity and/or specificity might have been evolutionarily relevant. To this end, we investigated the physiological context of the different clade IF enzymes characterized in our study. IF-1 and IF-2 are native to *Gaiellales* bacteria, which belong to the *Actinomycetoa*, a diverse phylum of Gram-positive bacteria^40^. IF-3 and IF-4 are found in representatives of the phylum *Chloroflexota*, which is comprised of green non-sulfur bacteria, some of which are aerobic thermophiles^41^. IF-5 is from an organism belonging to the class of *Limnochordia*, a not well-characterized class of moderately thermophilic bacteria of the *Firmicutes* phylum^42^. Metagenomic analysis identified a phosphoribulokinase homolog in all cases, indicating an active Calvin-Benson-Bassham cycle in these organisms. Notably, however, none of the host genomes contain genes that are essential for photosystem I or II, supporting the hypothesis that emergence of the SSU in Form I was independent of (and likely prior to) the evolution of oxygenic photosynthesis^18^.

To understand the relationship of the five characterized IF clade Rubiscos with an aerobic lifestyle or aerobic growth conditions, we further searched for the presence of oxygen-sensitive proteins or enzymes conferring oxygen tolerance in the respective genomes (Figure 3 B). The host of IF-4, IF-5 and IF-3 encode multiple oxygen-sensitive enzymes, in particular homologs of carbon monoxide dehydrogenase (IF-4 & IF-5), as well as pyruvate synthase (pyruvate:ferredoxin oxidoreductase, PFOR) or 2-oxoacid:acceptor oxidoreductase homologs (IF-3, IF-4, IF-5), suggesting a (facultative) anaerobic lifestyle for these organisms. Most hosts also encode a superoxide dismutase (SOD), while the host of IF-1 additionally encodes a catalase (CAT) homolog, indicating that these organisms are (at least) adapted to oxygen exposure.

The hosts of IF-3, IF-4 and IF-5 are putative (hyper)thermophiles^41,42^. Thus, we also investigated the activity and stability of the Form IF Rubiscos across a range of temperatures (Figure 3 D - H). Indeed, these homologs exhibit an increase in catalytic rate at temperatures up to 55 °C (IF-3 & IF-4) or even 70 °C (IF-5). In contrast, the activity of IF-1 and IF-2 peaked at 40 °C, consistent with their mesophilic context. In summary, these data provided additional insights into the evolutionary history and environmental context of diverse homologs from the early branching Form IF clade. These enzyme complexes probably evolved in (facultative) anaerobic, non-photosynthetic bacteria, in which discrimination against oxygen became only relevant later^18^.

## Conclusion

In this work, we investigated several representatives of the newly discovered Form IF clade of Rubisco. We show that proteins of this clade are highly active and assemble into active LSU-SSU complexes without the need for specific chaperones. These findings are unexpected considering the phylogenetic proximity of the IF clade to the IA/B clade, which are organized as “Rubiscosomes”, i.e., require multiple enzymes for production and functional operation of Rubisco (Figure 1).

Notably, IF-1 and IF-2 could be purified in the absence of their cognate SSUs. Both IF-1 and IF-2 LSU-only complexes show high activity (∼ 4 s^−1^), while maintaining specificities of CO_2_ over O_2_ of up to S_c/o_ of 40. This remarkably high catalytic performance of IF-1 and IF-2 without their native SSU represents a significant difference from any other known Form I Rubisco, where the absence of SSUs typically results in failure to assemble or dramatic loss of function^25,43^.

Yet even though the SSU is not needed for basic catalysis of the IF-1 and IF-2 core complexes, it is still able to further tune their catalytic properties, increasing their activity by a factor of three and further enhancing specificity. These findings resembles the situation of an ancestrally reconstructed Form I Rubisco that showed how an intrinsic allosteric network can be modulated by the SSU for altered kinetic parameters^25^. However, while in the latter case, activity was apparently traded for specificity, in case of IF-1 and IF-2, interaction with the SSU improves both parameters (specificity and activity) in parallel indicating that similar effects might be possible for further efforts in engineering Rubisco SSU and LSU.

During evolution, Rubiscos have become dependent on the SSU for full functionality. This entrenchment is uniformly observed across all high specificity Form I clades and affects the enzyme in multiple ways, including improved stability, solubility and catalysis^25^. In case of IF-1 and IF-2, the SSU increases catalytic activity and specificity, providing yet another example of how the SSU has become indispensable for proper function of the enzyme in its physiological context. Interestingly, IF-1 and IF-2 have escaped the notorious dependency of Form I Rubiscos on the SSU. As other Form IF Rubiscos do not form a stable and soluble LSU-only complex^43^, this feature must therefore have been specifically regained along the branch leading towards IF-1 and IF-2, further supported by incapability of even their closest neighbor IF-5 of forming a functional LSU-only protein (Figure S 1 D). It is very surprising that the two catalytically best performing enzymes in the clade have a less tightly bound subunit and show activity as LSU-only constructs. Most likely, the LSU-only construct is never actually formed *in vivo,* but the enzymes recruited mutations that trade a tightly bound SSU for a higher overall catalytic activity. This could be explained by reduced selective pressure for the preference of CO_2_ over O_2_, which enabled the Rubiscos of these organisms to explore a wider range within the mutational space whilst not being forced to maintain tight binding of the SSU. To our knowledge, this finding represents the only case of Form I Rubisco being able to potentially overcome its own solubility entrenchment, suggesting that the dependency on the SSU could be easier to overcome than generally assumed.

While our findings showcase a dramatic flexibility and dynamic across the evolutionary landscape of Rubisco, and the LSU-SSU interface in particular, we note that the current state in IF-1 and IF-2 might be semi-stable, as single mutations can quickly result in solubility entrenchments in the absence of the SSU (Figure 3 C). Nevertheless, the discovery of the chaperone-free and SSU-independent nature of clade IF Rubiscos provides new opportunities for the expression and engineering of Form I Rubiscos *in vitro* and, more critically, *in vivo*. Altogether, Form IF enzymes offer a promising foundation for future efforts to engineer and tailor Rubiscos with less biochemical and/or biophysical constrains, especially bypassing the need for additional subunits and chaperons for producing highly active and specific Rubiscos.

## Methods

Unless otherwise specified, chemicals were purchased from Sigma-Aldrich and Carl Roth in the highest available purity and quality. NaH^14^CO_3_ and K^14^CN were obtained from Hartmann Analytics (Germany) and PerkinElmer (MA), respectively. RuBP was enzymatically synthesized according to published protocols^44^. To determine specificity constants, [1-^3^H] RuBP was enzymatically produced from D-[2-^3^H] glucose (PerkinElmer, MA) following established procedures^45^. The Rubisco inhibitor 2-carboxyarabinitol-1,5-bisphosphate (CABP) was previously synthesized in two batches using either non-radioactive or [¹⁴C]-labeled KCN, following published methods^44^. The concentration of the non-radioactive CABP batch was determined by titrating a Rubisco preparation of known active site concentration (quantified with [^14^C]-CABP) with varying amounts of cold CABP and subsequently used at the determined concentration.

### Molecular cloning and vector construction

Gene fragments were ordered from Twist Bioscience. All primers were ordered from Eurofins. A list of all primers used in this study is provided in Supplementary Table 1. Genes encoding the Form IF proteins were designed to have a C-terminal His_6_-tag on the SSU and were cloned into the standard expression vector pET28b (Merck Chemicals) for IF Rubiscos and a pET16b for the IF-1 and IF-2 LSU only Rubisco. For LSU-SSU co-expression, a multicistronic cassette was constructed using an RBS encoding overhang sequence between the LSU and SSU as described in Schulz et al.^18^ Successful assembly of plasmids was verified by DNA sequencing (Microsynth).

Single-site mutants were created using an adapted Q5 SiteDirected Mutagenesis procedure. Mutagenesis primers were designed using NEBasechanger_v1 (nebasechangerv1.neb.com) and used to amplify the desired vector with Phusion High-Fidelity PCR Master Mix (Supplementary Table 1). Resulting PCR products were used in KLD enzyme mix (New England Biolabs) reactions and subsequently transformed before the resulting vector was purified and mutagenesis success was verified by sequencing.

### Protein Purification

For heterologous overexpression of genes encoding the Form IF proteins, the expression vector was transformed into chemically competent *E. coli* BL21 (DE3) cells (Thermo Fisher) and grown overnight on LB agar plates containing 50 µM/mL kanamycin (50 µM/mL Carbenicillin for IF-1 and IF-2 LSU only) at 37°C.

Colonies were used to inoculate an overnight preculture in LB media containing the respective antibiotics. For protein production, terrific broth (TB) media was inoculated using the pre-culture and grown over night at 25°C. Cells were harvested by centrifugation using a Beckmann centrifuge at 5,000 xg, 10 °C for 10 min and cell pellets were directly used for purification. Cell pellets were resuspended in buffer A (50 mM HEPES-NaOH, 500 mM NaCl, pH 7.6) and lysed using homogenizer (Avestin) with 4 cycles at 80 PSI. Lysate was clarified by centrifugation at 11,000 xg, at 15 °C for 1 hour. The resulting supernatant was filtered using 0.45 µm syringe tip filters, loaded on pre-equilibrated Protino Ni-NTA Agarose beads (Macherey-Nagel) in a gravity column, and washed using 10 column volumes (CV) buffer A. The protein-bound Ni-NTA resin was washed with 10 CV of 15% (v/v) buffer B (50 mM HEPES, 500 mM NaCl, 500 mM imidazole, pH 7.6) in buffer A and eluted in 4 CV buffer B. The eluate was concentrated using 50 kDa MWCO centrifugal filters (Amicon) to a total volume of 2.5 mL and either desalted using PD-10 desalting columns and desalting buffer (25 mM Tricine-NaOH, 75 mM NaCl, pH 8.0) for further experiments or for structure resolution further purified on a size-exclusion chromatography (SEC) on a Superdex HiLoad16/600 column. For SEC, the column was pre-equilibrated with desalting buffer and operated at a constant flow of 1 mL/min using an ÄKTA Pure system (GE Healthcare). Fractions with high UV-vis absorbance at 280 nm were analyzed using SDS-PAGE on 4 – 20 % gradient gels (Bio-Rad). Purified Rubiscos were either used immediately for experiments and crystallization (stored at 4 °C over night) or flash frozen in liquid nitrogen and further stored at – 70 °C.

### Solubility testing

For testing to introduce SSU independency in Form IF-5 Rubisco, samples were inoculated at an OD_600_ of 0.1 and incubated for 4 h at 37 °C. After induction with 1 mM IPTG, cultures were grown for 20 h at 25 °C. Cells were harvested in at 4000 xg for 5 min at 15 °C and resuspended in Lysis buffer (B-PER, thermo scientific), containing a spatula tip DNase, 1 mM PMSF and 0.2 mg mL^−1^ Lysozyme. Samples were incubated at 1000 rpm, 37 °C for 15 min. Lysate was clarified by centrifugation at 4000 xg, 15 °C for 15 min. Pellet and supernatant samples were collected for SDS PAGE.

### SDS-PAGE

For SDS-PAGE analysis, NuPAGE™ LDS Sample Buffer (4X) (Novex) was used to denature samples at 95 °C for 5 min. Denatured samples were loaded on 4 – 20 % Mini-PROTEAN TGX precast gradient gels (Bio-Rad) and protein separated by electrophoresis at 100 V for 40 min. Gel was stained using ReadyBlue Protein Gel Stain (Sigma).

### Kinetic measurements

^14^CO_2_ fixation assays were carried out in 7.7 mL septum-capped glass scintillation vials with a total volume of 500 µL. The assay buffer (100 mM EPPS-NaOH, 20 mM MgCl_2_, 1mM EDTA, pH 8.0) and further components were pre equilibrated with CO_2_ free N_2_ gas. For activations, carbonic anhydrase (0.01 mg mL^−1^), self-synthesized RuBP (1.08 mM), and NaH^14^CO_3_ were added. Dissolved CO_2_ concentrations (0 µM – 750 µM) were calculated using the Henderson-Hasselbalch equation with p*K* values for carbonic acid of pKa1 = 6.25 and pKa2 = 10.33, taking in account the assay and headspace volume 53 (Supplementary Table 2).

To quantify the active site of each Rubisco a [^14^C]-2-CABP binding assay was conducted on 0.4 µM purified Rubisco (monomer, as determined by UV-Vis measurements). Unbound [^14^C]-2-CABP was separated from CABP-bound Rubisco using 10 mL Sephadex G50 Fine resin in a glass chromatography column (Biorad Econo Column #7370732). Quantification of active sites was done by scintillation counting and calculated as described previously.^54^

Purified co-expressed LSU-SSU Rubisco was activated in assay buffer in the presence of 50 mM NaHCO_3_. For further investigation of IF-1_LS_ and IF-2_LS_ we also tested IF-1_L_ and IF-2_L_ and in presence of a tenfold excess of IF-1 SSU and IF-2 SSU respectively.

For reaction initiation, 20 µL activated Rubisco with final concentration of 0.4 µM for IF Rubiscos, was added to the reaction mixture and stopped after 2 min by quenching with 200 µL of 50% (v/v) formic acid. For LSU only and LSU supplemented with tenfold molar excess SSU, 40 µL activated Rubisco with a final concentration of 0.6 µM was used. Samples were stored at 95 °C overnight to ensure evaporation of the whole sample to quantify the remaining radioactivity after having all the CO_2_ gas out. Residual material was resuspended in 500 μL ddH2O, transferred to 7 mL scintillation vials, and mixed with 4.5 mL scintillation cocktail (ROTISZINT®HighCapacity). The amount of acid-stable radioactivity was quantified using a Beckman LS 6000 scintillation counter and used to determine the rate of carboxylation, after correcting for the scintillation counter efficiency (cpm/dpm) and background radioactivity (determined by running a no-RuBP control). The specific activity of the NaH^14^CO_3_ stock was measured by completely turning over 22.6 nM of RuBP using the highest employed NaH^14^CO_3_ concentration in a 60-minute reaction.

### Specificity Assay

Measurements for the specificity S_C/O_ for CO_2_ over O_2_, carbonic anhydrase and assay buffer (30 mM triethanolamine, 15 mM MgCl_2_, pH 8.3) were flushed with CO_2_-free N_2_ for at least one hour using 20 mL scintillation vials and then with 700 ppm CO_2_ in O_2_ for two hours. Purified Rubisco was activated by transferring it into the reaction mixture containing 0.02 mg mL^−1^ carbonic anhydrase at 25 °C for one hour and afterwards the reaction was initiated by adding [1H]^3^H-RuBP and stopped after running to completion after 2 hours by dephosphorylating with alkaline phosphatase (20 U/ reaction). Phosphoglycerate and phosphoglycolate were separated on an HPX-87H column (Bio-Rad) and the peaks were quantified using liquid scintillation counting. The S_C/O_ was subsequently calculated as described previously^18^.

### Specificity Assay for IF-1 10x SSU

Specificity assay was adapted from Schulz et al.^18^. Purified Rubisco was activated by transferring it into the reaction mixture containing 0.02 mg/mL carbonic anhydrase at 25 °C for one hour and afterwards the reaction was initiated by adding self-synthesized RuBP and stopped after running to completion after 3 hours by dephosphorylating with alkaline phosphatase (20 U/ reaction). Samples were centrifuged for 30 min at 10.000 xg and submitted for LC-MS/MS measurements. The S_C/O_ was subsequently calculated as described previously.^18,55^

### LC-MS/MS

For quantitative determination of produced glycerate and glycolate, LC-MS/MS was performed^55^. The chromatographic separation was performed using an Agilent Infinity II 1290 HPLC system using a Kinetex EVO C18 column (150 × 2.1 mm, 3 μm particle size, 100 Å pore size, Phenomenex) connected to a guard column of similar specificity (20 × 2.1 mm, 3 μm particle size, Phenomoenex) at a constant flow rate of 0.2 mL/min with mobile phase A being 0.1 % formic acid in water and phase B being 0.2 % formic acid in methanol (Honeywell, Morristown, New Jersey, USA) at 25 °C, using an injection volume of 1 µL. The profile of the mobile phase consisted of the following steps and linear gradients: 0 – 5 min constant at 0 % B; 5 – 6 min from 0 % to 100% B; 6 – 8 min constant at 100 % B; 8 – 8.1 min from 100 % to 0% B; 8.1 – 12 min constant at 0 % B. An Agilent 6495 mass spectrometer was used in negative mode with an electrospray ionization source and the following conditions: ESI spray voltage 2000 V, nozzle voltage 500 V, sheath gas 260 °C at 10 L/min, nebulizer pressure 35 psig and drying gas 100 °C at 13 L/min. Compounds were identified based on their mass transition and retention time compared to standards. Chromatograms were integrated using MassHunter software (Agilent, Santa Clara, CA, USA). Absolute concentrations were determined based on an external Standard curve.

### Heat Assay

Rubisco catalysis at elevated temperatures was determined using ^14^CO_2_ incorporation assays. Reactions were performed with NaH^14^CO_3_ stocks with specific activities between 45 – 135 mCi/μmol (corresponding to ∼100 – 300 cpm/nmol). Values for the solubility of CO_2_ in water (mol L^−1^ atm^−1^), the p*K*_a_1 and the p*K*_a_2 for NaHCO_3_ as well as the p*K*_a_ of the used buffer at elevated temperatures were taken from literature or extrapolated from literature values. Henderson-Hasselbalch equation parameters are listed in Supplementary Table 2 and were used to calculate dissolved CO_2_ concentrations using the Henderson-Hasselbalch equation, while accounting for the changes in temperature. 500 μL assays in 1.5 mL microcentrifuge tubes at 27, 40, 54, 68.5 and 79 °C contained 46.5, 54, 60.3, 65.5 and 68.3 mM NaH^14^CO_3_, respectively, resulting in a final dissolved CO_2_ concentration of ∼750 μM for all conditions. For catalysis, 20 μM Rubisco was activated in reactions containing 50 mM HEPES-NaOH (pH 8.0), 10 mM MgCl_2_, 20 mM NaHCO_3_ and 0.02 mg/mL carbonic anhydrase. Activation mixture was then diluted 10-fold into reaction mixture containing 100 mM HEPES (pH ∼8.0 – 8.3 at the respective temperatures), 20 mM MgCl_2_, indicated amounts of NaH^14^CO_3_ and 0.02 mg/mL carbonic anhydrase. After reaching the desired assay temperature, reactions were incubated for 2 min, before catalysis was initiated by the addition of 2 mM RuBP. Reactions were quenched using 10% v/v (final concentration) formic acid. Reaction tubes were opened and left to evaporate on a heating block at 95 °C. Residual material was re-suspended in 500 μL ddH_2_O, transferred to 7 mL scintillation vials, and mixed with 4.5 mL scintillation cocktail (ROTISZINT®HighCapacity). The amount of acid-stable radioactivity was quantified using a Beckman LS 6000 scintillation counter and used to determine the rate of carboxylation, after correcting for the scintillation counter efficiency (cpm/dpm) and background radioactivity (determined by running a no-Rubisco control). Activities at 25 °C were set to 100 %.

### Crystallography

To solve structures of CABP-bound Form IF Rubiscos, purified proteins were stored at 4 °C over night after running SEC. On the next day, the purified Rubiscos were incubated at 25 °C with a 2-fold molar excess of CABP (dissolved in 100 mM Bicine, 17.6 mM MgCl2, pH 8), stored for 1 hour at 37°C in the presence of 3 % CO_2_ and directly used for crystal plates.

Sitting-drop vapour-diffusion method was used for crystallization at 16 °C.

Form IF-2 (5 mg mL^−1^) was mixed 1:1 (final drop volume 1.2 µL) with 0.2 M Sodium acetate, 0.1 M Sodium citrate, pH 5.5, 10 %(w/v) PEG 4000. The reservoir was filled with 45 µL of solution B3 Screen. Crystals appeared within 2 days and soaked after 5 days supplemented with solution B and 27 % PEG 200 as Cryoprotection before freezing with liquid nitrogen.

X-ray diffraction data (Supplementary Table 3) were obtained at the P14 beamline of DESY (Hamburg, Germany). Data processing was carried out using the XDS^46^ and CCP4 (v.7.0)^47^ software packages. Structures were determined via molecular replacement with Phaser from the Phenix software package (v.1.21.2)^48^, constructed using Phenix.Autobuild, and refined using Phenix.Refine. Additional modeling, manual refinement, and ligand fitting were performed in Coot (v.0.9.8.3)^49^. Final positional and B-factor refinements, as well as water picking, were completed using Phenix.Refine. Figures were made using Pymol 2.5 (The PyMOL Molecular Graphics System, Version 2.5 Schrödinger, LLC (www.pymol.org)).

The final Rubisco structure was deposited in the Protein Data Bank under accession number PDB 28VS (IF-2_LS_).

### Cryo-EM

For preparing IF-1_LS_, IF-2 _LS_ and IF-2 _L_ Cyro-EM grids, 3 µL Protein (1 mg mL^−1^) in desalting buffer (25 mM Tricine, pH 8.0, 75 mM NaCl) was supplemented with a twofold molar excess of CABP and 1.33 mM MgCl_2_ and activated for 20 min at 37 °C at 4 % CO_2_. Activated Rubisco was applied to R flat 2/1 Cu 200 mesh grids (QuantiFoil), which were glow-discharged for 45 s immediately before use. Grids were blotted for 9 s at blot force - 2 and plunge-frozen in liquid ethane cooled by liquid nitrogen using a Vitrobot Mark IV (Thermo Scientific).

Cryo-EM Data (Supplementary Table 4) were collected on a Titan Krios G3i transmission electron microscope (Thermo Scientific) operated at 300 kV, equipped with a BioQuantum energy filter and a K3 direct electron detector (Gatan). Electron counting super-resolution mode was performed at a nominal magnification of 105,000x (0.837 Å / 0.827 Å per pixel) with a total dose of 55 e^−^/Å^2^, using the aberration-free image-shift (AFIS) correction in EPU (Thermo Scientific). Five images were acquired per foil hole with nominal defocus range used for data collection was -0.8 to -2.0 μm. All image processing was performed in CryoSPARC (Structura Biotechnology Inc., Canada, v4.6)^50^. Dose-fractionated movies were gain-normalized, aligned, and dose-weighted using Patch Motion Correction, and contrast transfer function (CTF) parameters were estimated with Patch CTF. In total, 1,428,008 (IF-1_LS_), 200,583 (IF-2_L_), 262,072 (IF-2_LS_) candidate particles were extracted using a box size of 300 (IF-1_LS_), 600 (IF-2_L_), 380 (IF-2_LS_) pixels and subjected to repeated 2D classification to remove non-particle images. Cleaned particles were used for *ab initio* reconstruction using up to 6 classes, of which the best-resolved class was selected for further refinement.

The selected class was refined using non-uniform refinement with iterative per-particle and global tilt and trefoil aberration correction, yielding a final global resolution of 2.06 Å for IF-1_LS_ and 2.15 Å for IF-2_L_, 2.58 Å for IF-2_LS_ and a temperature factor of 74.7 (IF-1_LS_), 46.1 (IF-2_L_), 56.0 (IF-2_LS_) Å^2^.

Model interpretation was performed in COOT (v.0.9.8.3)^49^ using PDB entry 7QSW for IF-1_LS_, 7QSV for IF-2_L_ as an initial reference. The model was manually adjusted and iteratively refined by real-space refinement in Phenix (v. 1.21.2)^48^. Model quality was assessed using MolProbity^51^. Structural figures were generated using UCSF ChimeraX (v.1.7)^52^, and PyMOL (v.2.5). The final Rubisco filament structure and cryo-EM maps were deposited in the Protein Data Bank under accession number PDB 28VR (IF-1_LS_) and 28VT (IF-2 _L_) and in the Electron Microscopy Data Bank under accession EMDB 56893 (IF-1 _LS_), EMDB 56894 (IF-2 _L_), EMDB 56895 (IF-2_LS_ partially occupied) respectively.

### Mass Photometry

Microscope coverslips (1.5 H, 24 x 60 mm, Carl Roth) and CultureWell™ Reusable Gaskets (CW-50R-1.0, 50 wells, 3 mm diameter × 1 mm depth) were cleaned through three consecutive rinses with ddH_2_O and 100% isopropanol, followed by drying under a stream of pressurized air. For measurements, the gaskets were assembled on the coverslips and positioned on the stage of an OneMP mass photometer (MP, Refeyn Ltd, Oxford, UK) using immersion oil.

To prepare for measurements, the gasket wells were filled with 18 μL of 1x phosphate-buffered saline (PBS, 10 mM Na_2_HPO_4_, 1.8 mM KH_2_PO_4,_137 mM NaCl, 2.7 mM KCl, pH 7.4) to focus the glass surface. After focusing, 2 μL of sample was added, rapidly mixed while maintaining the focal position, and data acquisition commenced within 15 seconds. Data was collected for 60 seconds at 100 frames per second using AcquireMP (Refeyn Ltd, version 1.2.1). Protein samples were prepared by diluting purified protein to a concentration of 10 μM (monomer equivalent) in desalting buffer (25 mM Tricine, pH 8.0, 75 mM NaCl), as determined by absorption at 280 nm. Immediately prior to measurements, samples were diluted to a final concentration of 500 – 1000 nM, and 2 μL of the diluted sample was mixed with 18 μL of PBS used for focusing the mass photometer (MP) on the glass slide. MP contrast values were calibrated to molecular masses using the NativeMark™ Unstained Protein Standard (Thermo Fisher). For calibration, a 30-fold dilution of the 1x NativeMark stock in 1x PBS was prepared and used as a 10x working stock (2 μL of standard added to 18 μL of 1x PBS). Calibration measurements revealed well-resolved peaks at contrast values corresponding to molecular masses of 66 kDa, 146 kDa, 480 kDa, and 1048 kDa. These peaks were integrated and fitted to a linear regression using DiscoverMP (Refeyn Ltd, version 1.2.3). All mass photometry images were processed and analyzed using DiscoverMP (Refeyn Ltd, version 1.2.3).

## Acknowledgements

F.O, H.W., J.Z., A.M.K, S.P., T.J.E., and G.K.A.H. are grateful for generous support from the Max Planck Society. A.M.K. is grateful for funding provided by EMBO (ALTF 684-2022) and MSCA (Project 101106795 ECOFix). The synchrotron data was collected at beamline P13 operated by EMBL Hamburg at the PETRA III storage ring (DESY, Hamburg, Germany). We would like to thank Michael Agthe for the assistance in using the beamline. We gratefully thank the European Research Council for the ERC Grant „pro2neo-RUBISCO“ Grant agreement ID: 101140565

## Author Contributions

F.O, H.W, L.S., T.J.E., and G.K.A.H. conceived the project, analyzed data, and planned experiments. L.S. performed molecular work and MP measurements. F.O. performed protein purification, biochemistry, and MP measurements, molecular work, cryo-EM and crystal structures, enzyme kinetic analysis and specificity constant measurements. H.W performed phylogenetics, metagenomic investigations, enzyme kinetics, and molecular work. K.P.F. performed molecular work, MP measurements, and specificity constant measurements. A.M.K. processed the cryo-EM datasets. J.Z. collected, solved, refined, and analyzed x-ray structures and refined the cryo-EM structure. S.P. performed cryo-EM grid preparation and collected cryo-EM datasets. A.M.K., P.C., and N.P. optimized, prepared and measured the LC-MS/MS methods and samples. T.J.E., and G.K.A.H. supervised the project. F.O., H.W, T.J.E., and G.K.A.H. wrote the manuscript with contributions and comments from all authors.

## Data availability

All atomic models were deposited into the PDB under the IDs 28VR, 28VS, and 28VT, respectively. Cryo-EM maps were deposited into the EMDB under the IDs 56893, 56894, and 56895, respectively (see Supplementary Table 3 & Supplementary Table 4).

## Supplementary Figures

**Figure S1.**
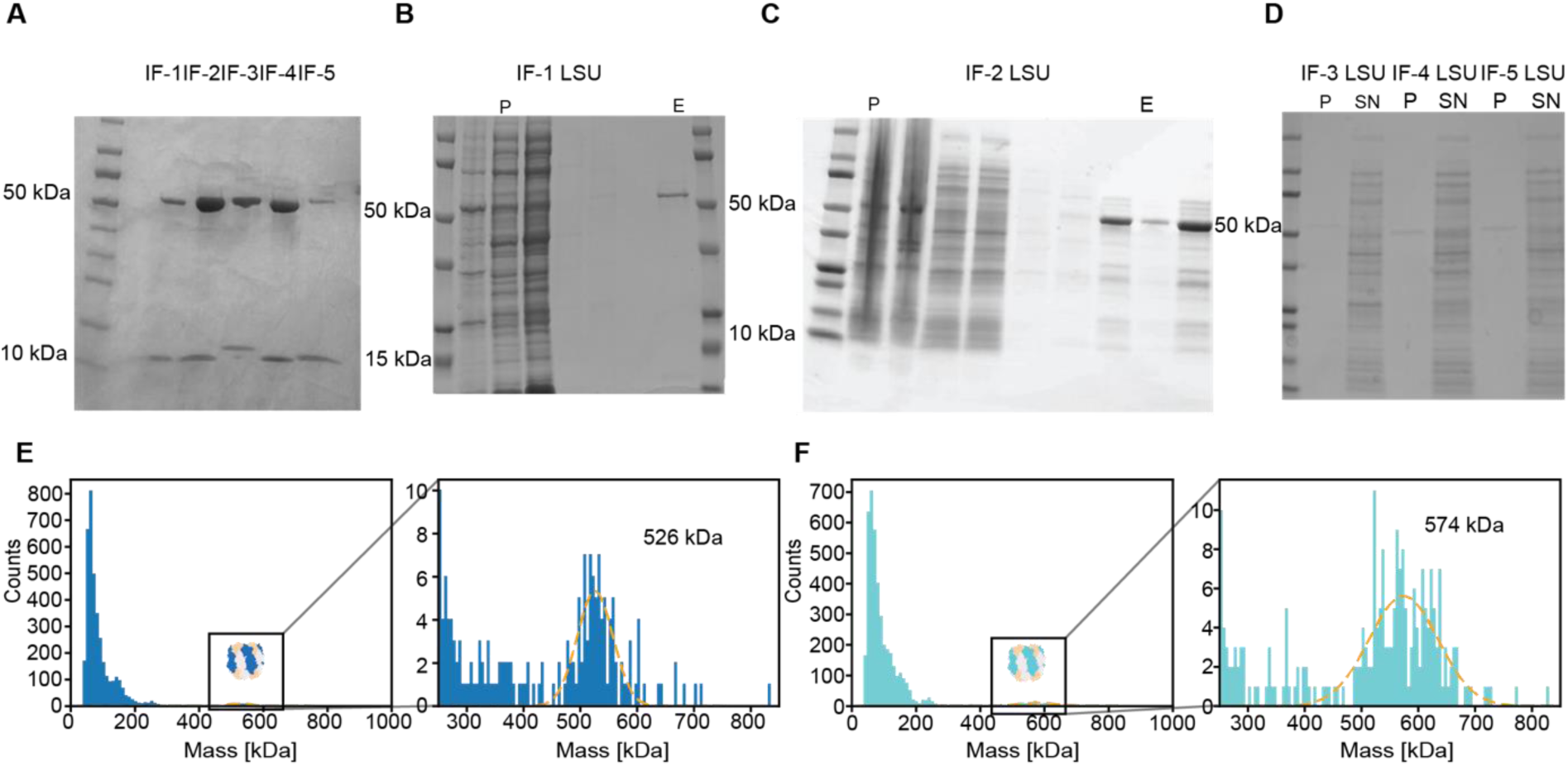
SDS PAGEs of Form IF Rubiscos and Form IF LSU only. Bands at 50 kDa indicates the LSU while bands above 10 kDa show the SSU. (**A**) Depicted are the co-expressed IF Rubiscos. (**B**), (**C**) For IF-1 and IF-2 an LSU only no SSU bands are observed. (**D**) IF-3, IF-4 and IF-5 show no soluble protein for the LSU. (**E**), (**F**) Showing the mass photometry of IF-1 and IF-2 in 2-fold molar excess of their respective small subunit.

**Figure S2.**
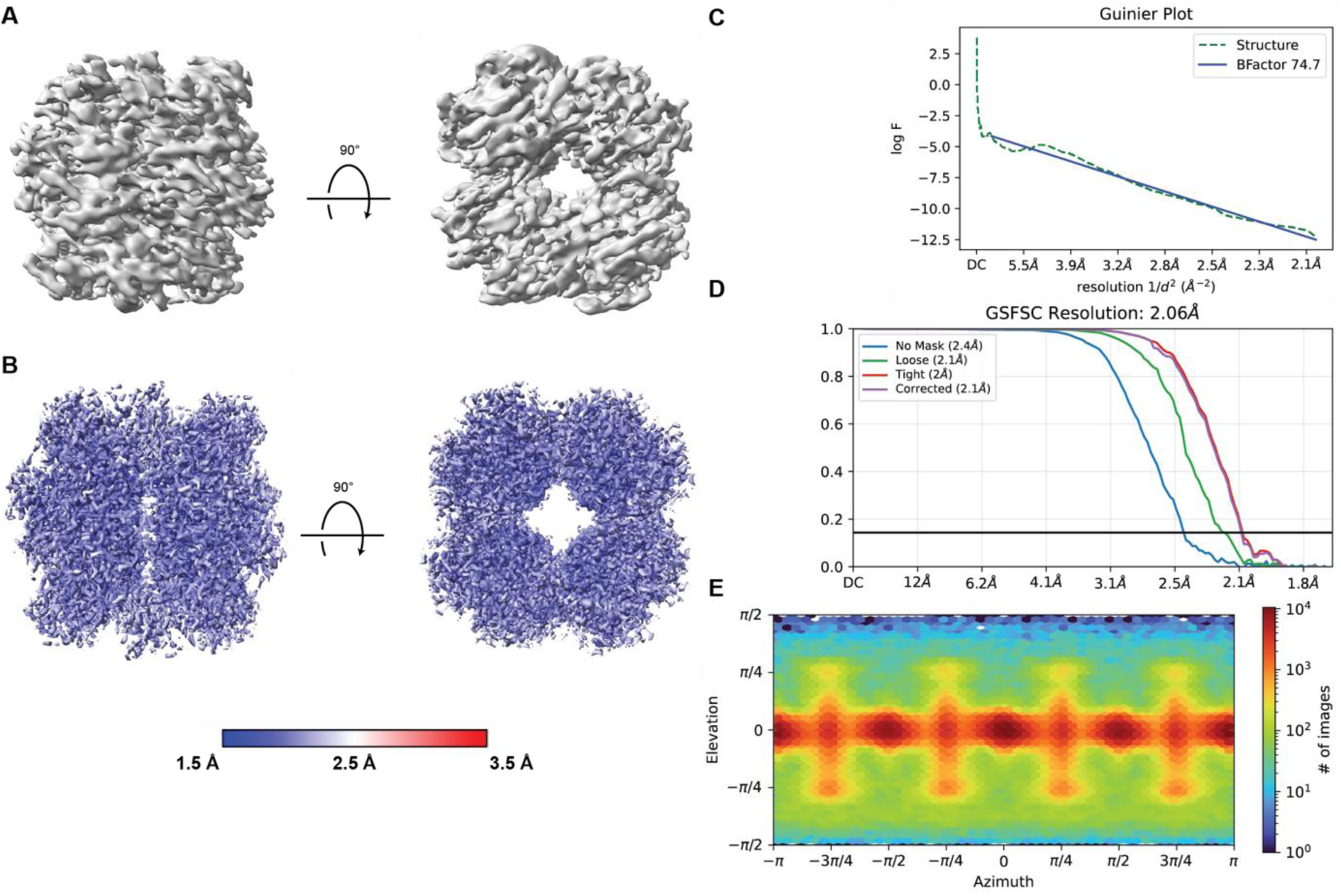
(**A**) Density map of the ab initio reconstructed volume. (**B**) Local estimated resolution map calculated by CryoSPARC mapped on the refined density of IF-1_LS_. Left side front view, right top view. (**C**) Guinier plot of the local refinement is shown indicating a B-factor of 74.7. (**D**) The GSFCS plot for IF-1_LS_ from the final local refinement is presented indicating a final global resolution of 2.06 Å. (**E**) Angular distribution of particles used for refinement.

**Figure S3.**
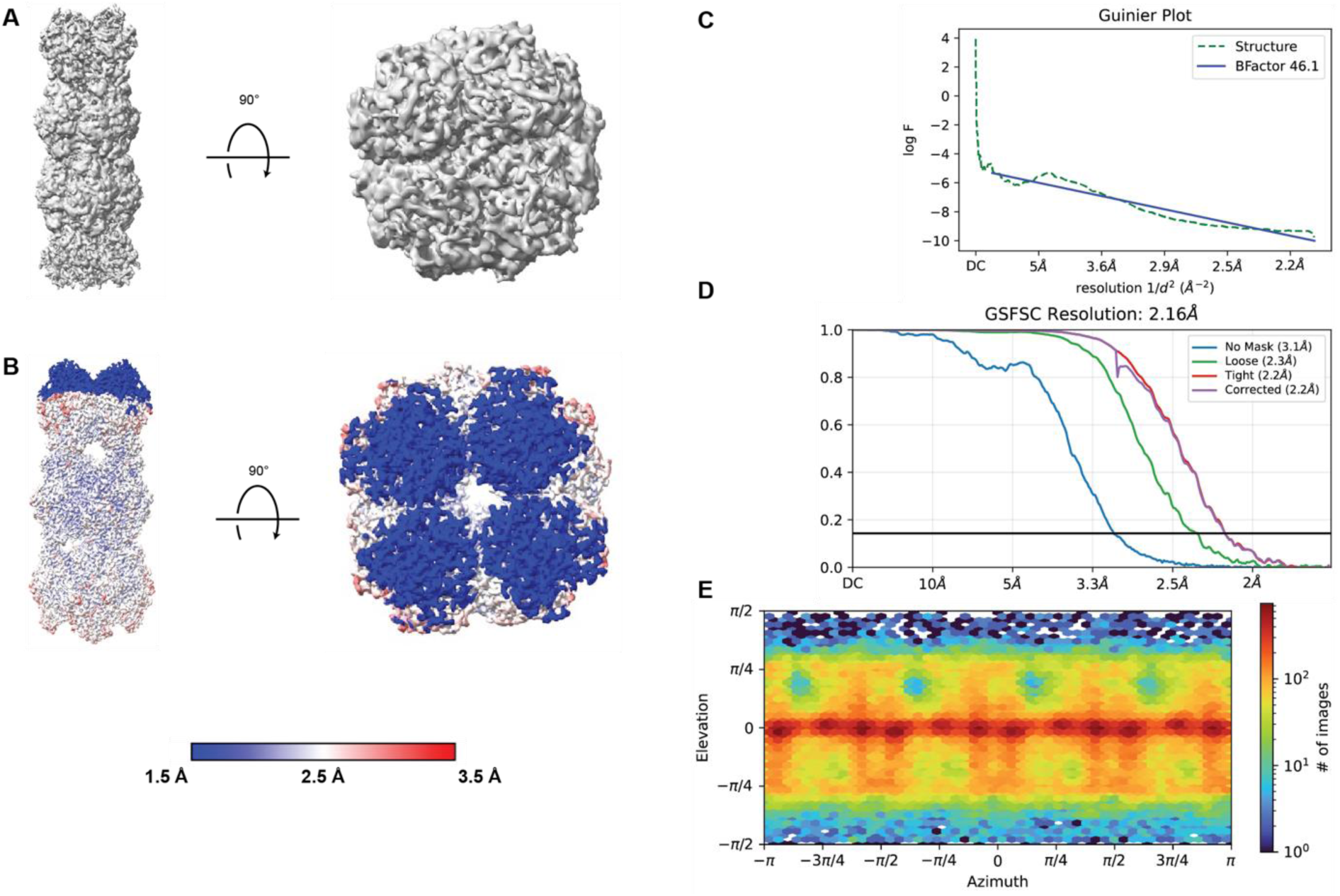
(**A**) Density map of the ab initio reconstructed volume. (**B**) Local estimated resolution map calculated by CryoSPARC mapped on the refined density of IF-2_L_. Left side front view, right top view of IF-2_L_. (**C**) Guinier plot of the local refinement is shown indicating a B-factor of 46.1. (**D**) The GSFCS plot for IF-2_L_ from the final local refinement is presented indicating a final global resolution of 2.16 Å. (**E**) Angular distribution of particles used for refinement.

**Figure S4.**
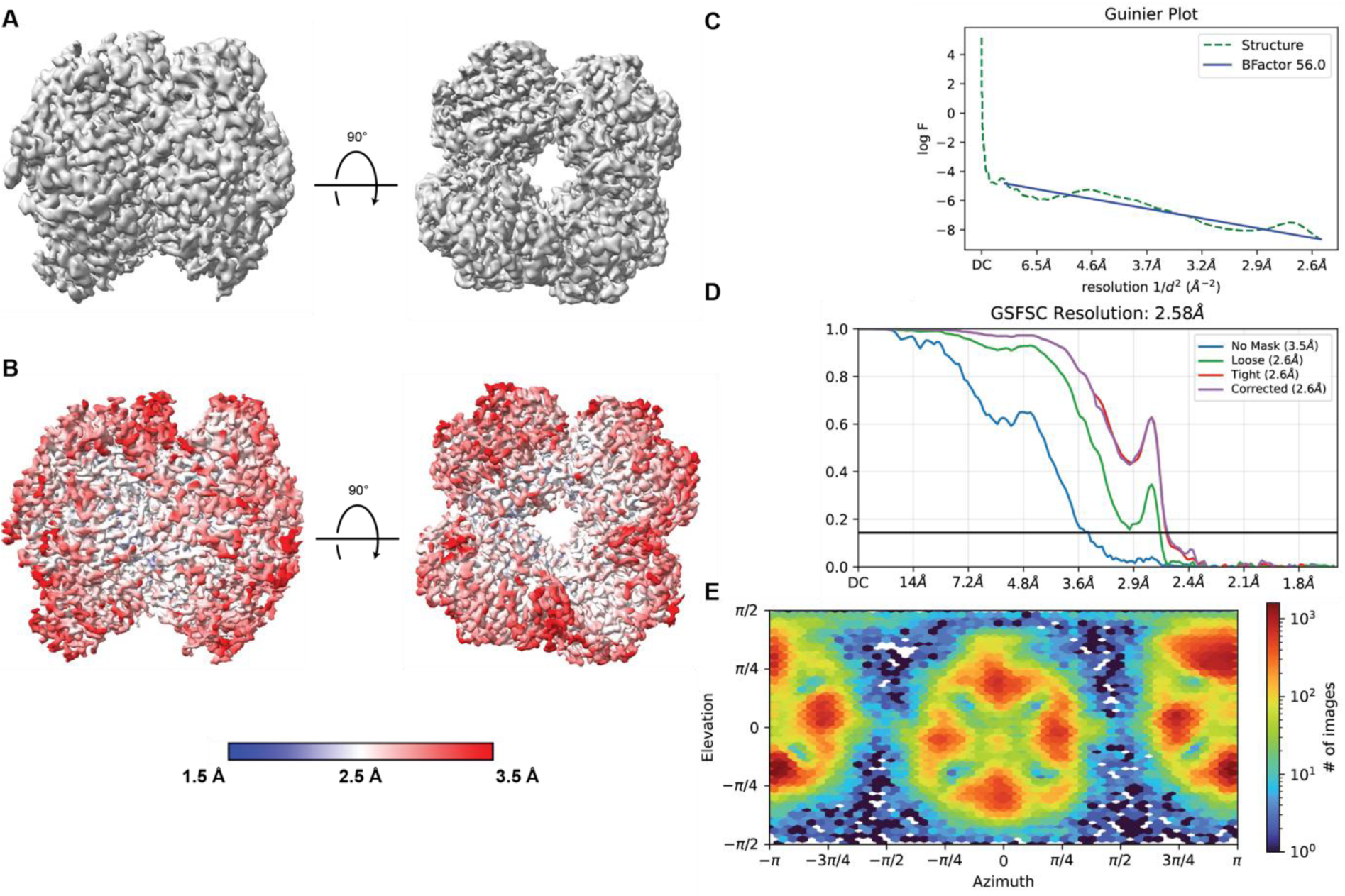
(**A**) Density map of the ab initio reconstructed volume. (**B**) Local estimated resolution map calculated by CryoSPARC mapped on the refined density of IF-2_LS_. Left side front view, right top view of IF-2_LS_. (**C**) Guinier plot of the local refinement is shown indicating a B-factor of 56.0. (**D**) The GSFCS plot for IF-2_LS_ from the final local refinement is presented indicating a final global resolution of 2.58 Å. (**E**) Angular distribution of particles used for refinement.

## Supplementary Tables

**Supplementary Table 1.**
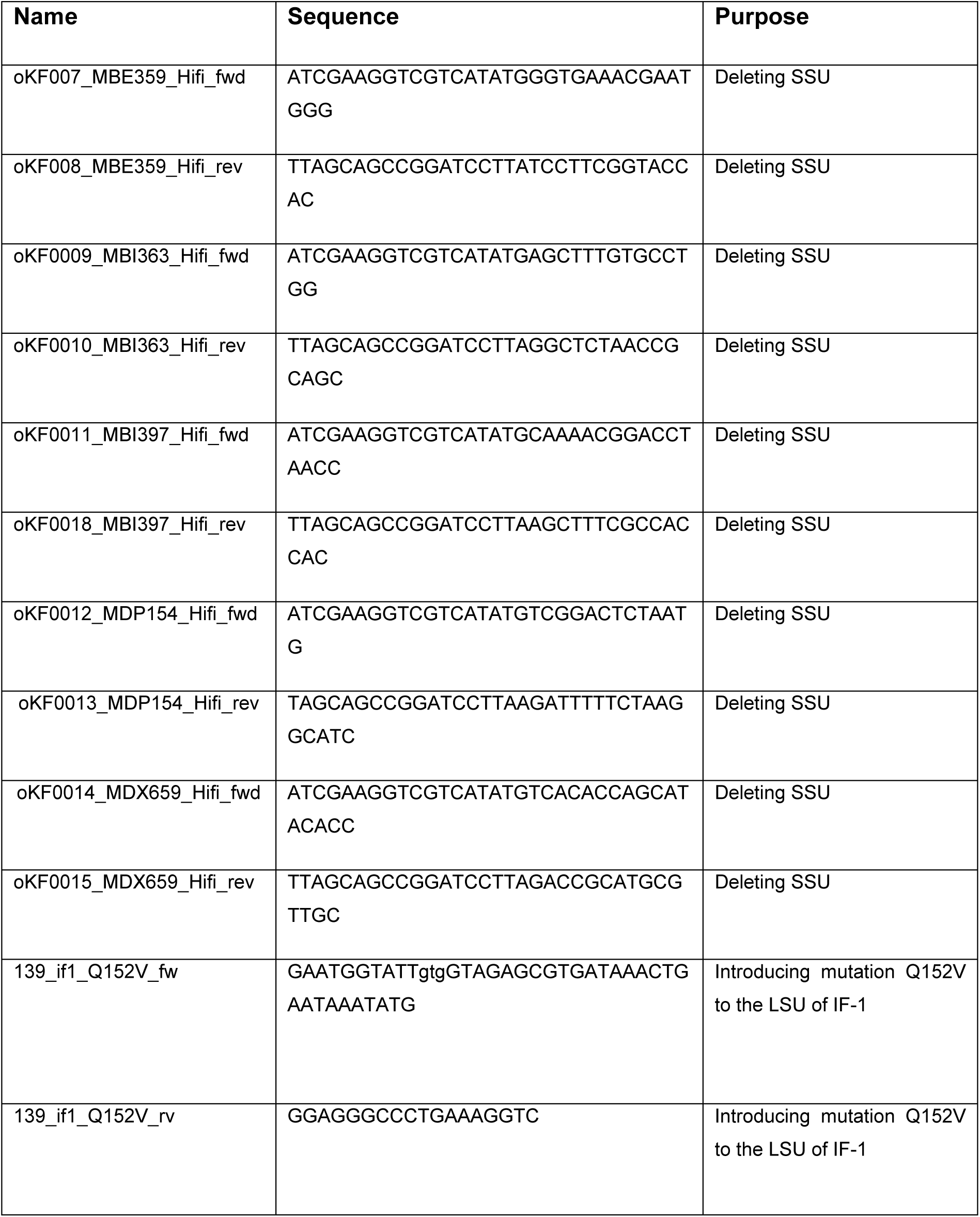
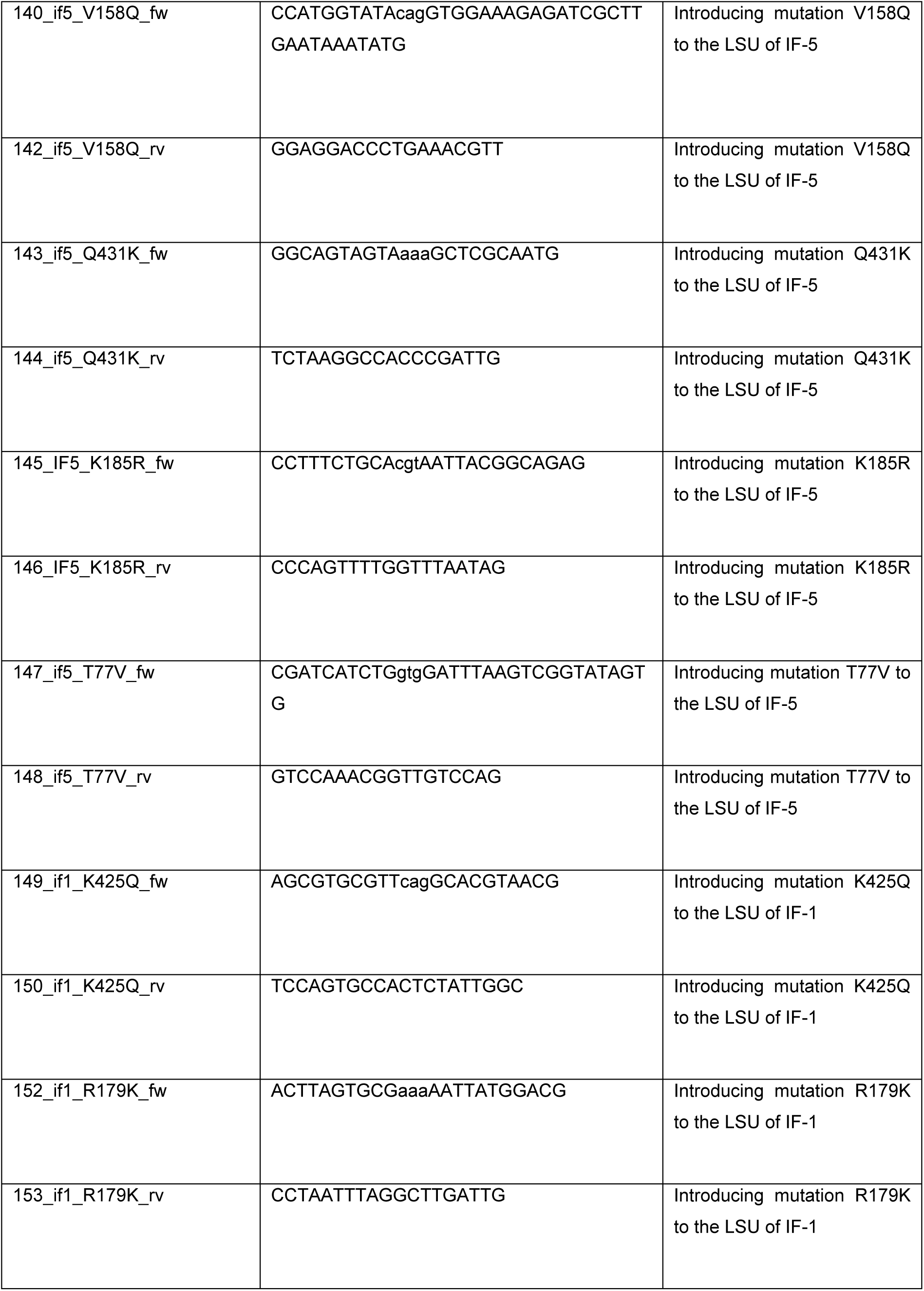
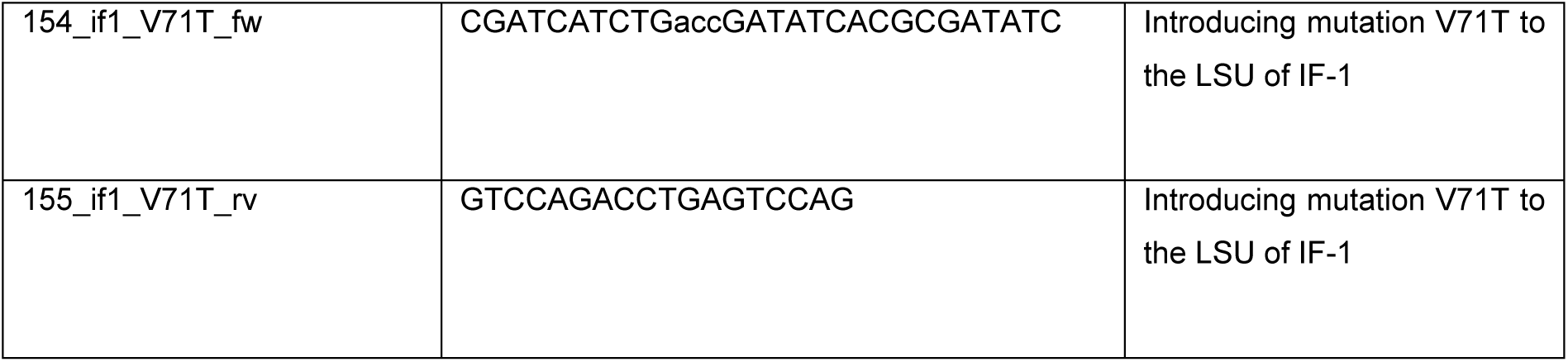
Primers used in this study.

**Supplementary Table 2.** Henderson-Hasselbalch coefficients at elevated temperatures. Solubility coefficient and pK_a_1/pK_a_2 of bicarbonate/CO_2_ for temperatures >50 °C (marked with an asterisk) were estimated from literature values at lower values (cited in source column) due to a lack of published values.

| Temperature | Solubility coefficient (Q) | $pK_{a1}$ ($H_2CO_3$) | $pK_{a2}$ ($H_2CO_3$) | Buffer pH at $25\text{ }^{\circ}C$ | source |
| --- | --- | --- | --- | --- | --- |
| $25\text{ }^{\circ}C$ | 0.03292 | 6,25 | 10,33 | 8 | 53,56,57 |
| $40\text{ }^{\circ}C$ | 0.02256 | 6,17 | 10,22 | 8,18 | 53,56,57 |
| $54\text{ }^{\circ}C$ * | 0.01581 | 6,13 | 10,15 | 8,37 | Estimated from literature |
| $69\text{ }^{\circ}C$ * | 0.01146 | 6,1 | 10,1 | 8,57 | Estimated from literature |
| $79\text{ }^{\circ}C$ * | 0.00943 | 6,086 | 10,07 | 8,72 | Estimated from literature |

**Supplementary Table 3.**
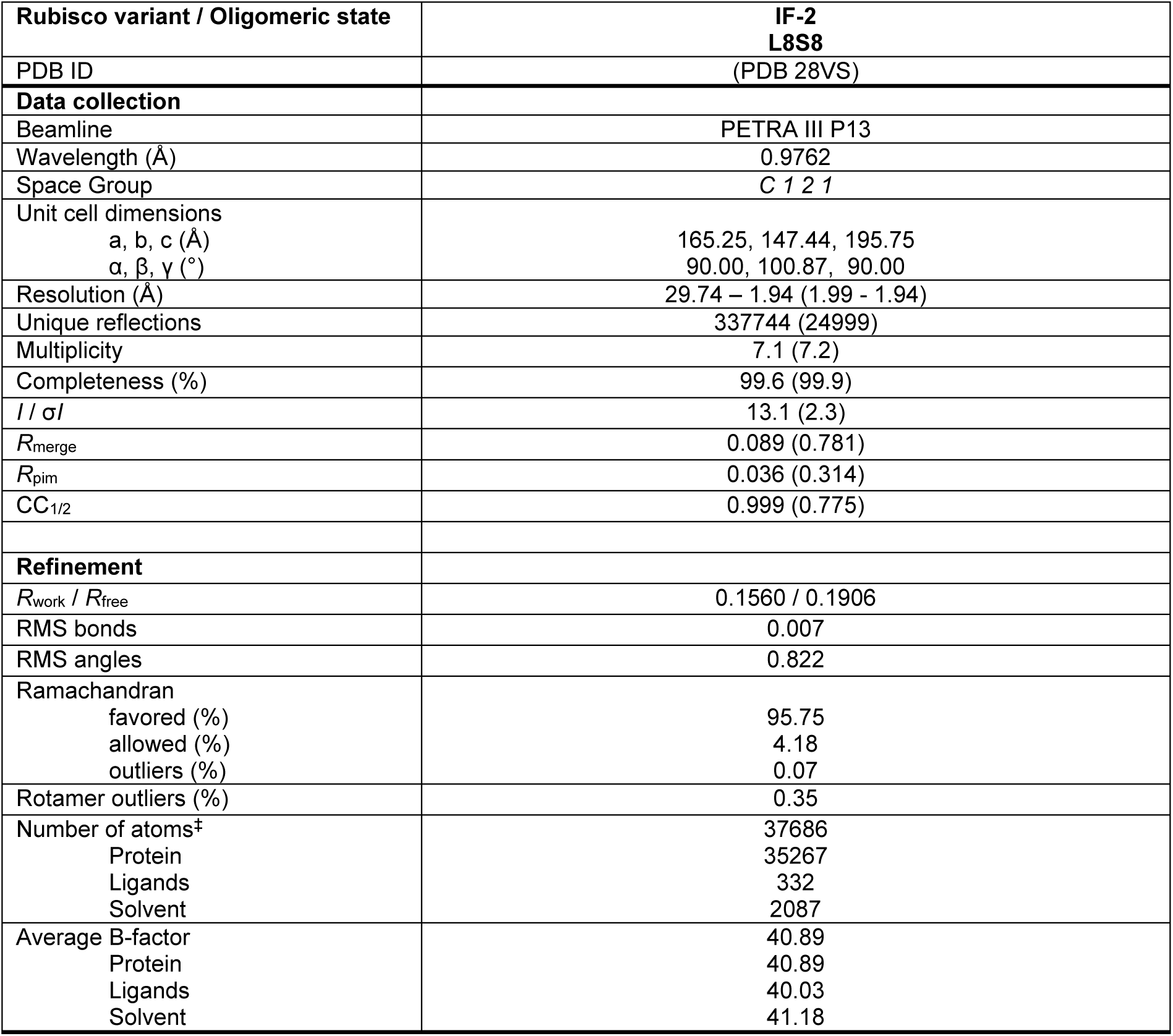
X-ray diffraction data collection and model refinement statistics. Values in parentheses are for highest-resolution shell.

| Rubisco variant / Oligomeric state | IF-2<br>L8S8 |
| --- | --- |
| PDB ID | (PDB 28VS) |
| **Data collection** |  |
| Beamline | PETRA III P13 |
| Wavelength (Å) | 0.9762 |
| Space Group | C 1 2 1 |
| Unit cell dimensions<br>a, b, c (Å)<br>$\alpha$, $\beta$, $\gamma$ (°) | 165.25, 147.44, 195.75<br>90.00, 100.87, 90.00 |
| Resolution (Å) | 29.74 – 1.94 (1.99 - 1.94) |
| Unique reflections | 337744 (24999) |
| Multiplicity | 7.1 (7.2) |
| Completeness (%) | 99.6 (99.9) |
| $I / \sigma I$ | 13.1 (2.3) |
| $R_{\text{merge}}$ | 0.089 (0.781) |
| $R_{\text{pim}}$ | 0.036 (0.314) |
| $CC_{1/2}$ | 0.999 (0.775) |
| **Refinement** |  |
| $R_{\text{work}} / R_{\text{free}}$ | 0.1560 / 0.1906 |
| RMS bonds | 0.007 |
| RMS angles | 0.822 |
| Ramachandran<br>favored (%)<br>allowed (%)<br>outliers (%) | 95.75<br>4.18<br>0.07 |
| Rotamer outliers (%) | 0.35 |
| Number of atoms<sup>‡</sup><br>Protein<br>Ligands<br>Solvent | 37686<br>35267<br>332<br>2087 |
| Average B-factor<br>Protein<br>Ligands<br>Solvent | 40.89<br>40.89<br>40.03<br>41.18 |

**Supplementary Table 4.** Cryo-EM data collection, image processing, and refinement.

| Rubisco variant / Oligomeric state | IF-1 / L8S8 | IF-2 / 3x L8 | IF-2 / L8S1 |
| --- | --- | --- | --- |
| Accession IDs | PDB 28VR, EMDB 56893 | PDB 28VT, EMDB 56894 | EMDB 56895 |
| Data collection |  |  |  |
| Microscope | Titan Krios G3i |  |  |
| Voltage (kV) | 300 |  |  |
| Camera | K3 |  |  |
| Magnification | 130,000 | 130,000 | 130,000 |
| Pixel size at detector (Å/pixel) | 0.827 | 0.837 | 0.827 |
| Total electron exposure (e<sup>-</sup>/Å<sup>2</sup>) | 55 | 55 | 55 |
| Frames per exposure | 55 | 55 | 55 |
| Defocus range (µm) | -0.8 to - 2.0 | -0.8 to - 2.0 | -0.8 to - 2.0 |
| Automation software | EPU, CryoSPARC |  |  |
| Total extracted particles (no.) | 4,289,980 | 2,555,801 | 818,461 |
| Final particles (no.) | 1,428,009 | 200,583 | 262,072 |
| Point-group | D4 | C4 | C1 |
| Resolution (global, Å) | 2.06 | 2.15 | 2.58 |
| FSC<sub>0.143</sub> (unmasked / masked) | (2.42 / 2.06) | (3.05 / 2.15) | (3.49 / 2.58) |
| Resolution range (local, Å) | 1.91-2.12 | 2.07-2.19 | 2.39-3.15 |
| Map sharpening B-factor (Å<sup>2</sup>) | 120.1 | 46.1 | 56.0 |
| Map sharpening methods | CryoSPARC, Phenix anisotropic | CryoSPARC |  |
| Model Refinement |  |  |  |
| Refinement package | Phenix |  |  |
| method | Real space |  |  |
| Resolution cutoff (Å) | 2.04 | 2.14 |  |
| Model-Map scores |  |  |  |
| CC<sub>volume</sub> | 0.87 | 0.89 |  |
| avg. FSC<sub>0.143</sub> (unmasked / masked) | 2.05 / 2.04 | 2.15 / 2.13 |  |
| Number of atoms | 35,618 | 87,896 |  |
| B-factors (Å<sup>2</sup>) |  |  |  |
| Protein (min / max / mean) | 1.94 / 74.14 / 17.87 | 14.07 / 122.24 / 46.10 |  |
| Ligands (min / max / mean) | 11.98 / 26.61 / 19.00 | 37.61 / 122.89 / 59.81 |  |
| Water (min / max / mean) | 4.81 / 24.73 / 14.87 | 20.95 / 45.46 / 34.63 |  |
| RMS angles | 0.651 | 0.517 |  |
| RMS bonds | 0.004 | 0.002 |  |
| Model Validation |  |  |  |
| Ramachandran |  |  |  |
| favored (%) | 96.80 | 97.93 |  |
| allowed (%) | 3.20 | 2.07 |  |
| outliers (%) | 0.00 | 0.00 |  |
| Rotamer outliers (%) | 0.17 | 0.44 |  |
| MolProbity score | 1.54 | 1.21 |  |
| CaBLAM outliers (%) | 1.99 | 1.34 |  |
| C-beta deviations | 0.00 | 0.00 |  |
| Clashscore | 6.26 | 4.08 |  |

## Sequences

Form IF-1 LSU:

ATGTCTGACCATCACACCCCGTTATTCGCAGCGGGTGTAAAAGAATACCGCGAGACTTACTGGAGACCTGGGTATGATCCTGCCGATACAGATATACTGACGGCTTTTCGTGTTACACCGCAGCCAGGTGTCCCGCTGGAAGAAGCAGCCGCCGCGGTCGCCGCGGAGAGCAGCACTGGTACATGGACCCAGGTCTGGACGGATCATCTTGTCGATATAGCACGGTACCAAGCTAAATGCTATCGTATAGAGCCGGTGCCGGGCGAGCCGGACCAGGCCATTGCGTTCATCGCATATCCTTTAGACCTCTTCGAAGAAGGATCGGTTAGTAATGTGATGTCCAGTATCGTTGGCAATGTATTTGGTTTTAAGGCTCTGCGCGCGCTCCGACTGGAAGATATGCGCATCCCGCCCGCTTACATGAAAACATTTCAGGGCCCGCCGAACGGTATCCAAGTTGAACGTGATAAACTCAATAAGTATGGTCGCCCAATGTTAGGTTGTACCATTAAACCCAAGCTGGGCCTGTCTGCCCGCAACTATGGTCGTGCAGTGTATGAAGCCCTGCGTGGTGGGCTTGACTTTACGAAGGATGATGAAAACATTAATTCACAGCCTTTTATGCGCTGGCGTGACAGATATGAATTCGTGATGGAGGCAGTGAAAAAAGCCGAAAGCGAAACGGGCGAACGTAAGGGCCACTATCTTAACGTAACAGCTCCCACAGTGGAACAAACCTTCGAGCGTGCGGAAGTTGCTAAACTTTTGGGTTCGCCTATTATAATGCATGATTACCTGACGGCGGGCTTTGCGATGCATCAGAGCTTGTCACACTGGTGCCGGGCGAACGGGATGTTATTACACGTGCATCGCGCAATGCACGCGGTAATGGACCGCCAAAAAAATCATGGCATTCATTTCCGAGTGCTGGCCAAATGGCTGCGAATGGCTGGGGGCGACCATTTACATGTGGGTACCGTCGTGGGTAAGTTAGAAGGTGATAGAGCTTCTACACTGGGTGTGATTGACCTCCTGCGCTTAGATAATGTGCCGGCCGACCCTTCCCGGGGCCTGTACTTCGATCAGCCCTGGGTGTCCCTCCCGGCTGTCATGCCGGTCGCGTCCGGCGGTATTCATGTCCATCATATGCCGGATTTAGTGGAAATATTTGGAGATGATTCTGTATTACAATTTGGTGGCGGCACTTTAGGGCACCCCTGGGGTAACGCGGCAGGGGCTACCGCGAACCGAGTAGCACTTGAAGCATGCGTCAAAGCTCGCAACGAAGGACGTGCGCTGATTGCGGAGGGACCTCAGATTTTACGTGAAGCCGCAGATCATTCACCGGAGCTGGCTGTTGCCATGGAGACATGGAAGGACATACGTTTCGAGCAGCCAACCGTTGATACTCTGGACGCCCAACATGCTGTTTAA

Form IF-1 SSU:

ATGCCATCCGAGACCTTTTCATATTTACCGACTCTCACCGATGCGGAGGTCCGTGCACAGGTGGCTAGTCTGGTGCGTGAAGGTCTGATTCCGGGGCTGGAATTCACCCATGATCCGTCCCCGCGTGACTCCTTTTGGAGCCTCTGGAAACTGCCGTTCGTAGAACGCCCGAGTCCAGACGCGGTCCTGGCGGAAATTGATCAGTGCGCGGCGGAACACCCGGACGCCCATA TCAAGCTTGTTGGATATGATTCCATACGCCAAGGCCAAGTGGTCGCATTTGTTGTTAGAGTTCCGCGC

Form IF-2 LSU:

ATGTCACACCAGCATACACCCCTGTTTGCTGCAGGCGTAAAAGAATACCGGGAAACTTACTGGCAGCCGGGCTATGAGCCCGCCGATACGGATATTTTAACCGCATTCCGTGTGACCCCACAGCCGGGGGTCCCAATCGAAGAAGCAGCAGCTGCAGTTGCTGCTGAGAGCTCAACCGGGACCTGGACTCAGGTCTGGACCGATCATCTGGTGGATATCACGCGATATCAGGCGAAATGTTATCGCCTGGCGCCAGTGCCAGCCGAACCGGATCAGGCGATTGCCTTTATAGCATATCCCCTTGATCTCTTTGAAGAAGGTTCTGTTAGTAATGTAATGTCTTCAATCGTTGGGAATGTCTTCGGCTTTAAAGCGCTGCGGGCATTGCGCCTGGAAGATATGCGAATCCCACCGGCATACATGAAGACCTTTCAGGGCCCTCCGAATGGTATTCAGGTAGAGCGTGATAAACTGAATAAATATGGTCGTCCAATGCTGGGGTGTACAATCAAGCCTAAATTAGGACTTAGTGCGCGTAATTATGGACGGGCGGTCTACGAAGCACTGCGTGGTGGTCTTGATTTCACGAAAGACGATGAGAATATAAACAGCCAACCATTTATGCGCTGGCGTGATCGTTATGAATTTGTGATGGAAGCGGTAAAAAAAGCTGAGAGCGAGACTGGCGAAAGAAAGGGTCACTATTTAAATGTGACTGCACCCACCGTTGAACAGACCTTTGAAAGAGCCGAAGTAGCTAAACAGTTAGGTGCACCTATCATCATGCATGATTACCTGACGGCTGGATTCGCAATGCATCAGTCGCTGTCACATTGGTGTCGTGCCAACGGTATGCTCTTACATGTTCATCGTGCAATGCATGCCGTTATGGATCGTCAAAAAAACCATGGCATCCACTTTCGTGTCTTAGCCAAATGGTTGCGCATGGCGGGCGGCGACCACCTGCATGTTGGTACGGTCGTGGGTAAACTGGAAGGCGATCGGGCCAGCACGCTTGGTGTTATCGACCTTCTGCGTCTTGACCACATACCAGCAGACCCTGCAAGAGGTTTATACTTTGACCAGCCATGGGTGTCCCTCCCGGCTGTGATGCCTGTAGCGAGTGGCGGGATTCATGTACATCACATGCCGGACCTGGTAGAAATTTTTGGCGACGACAGCGTTCTGCAATTTGGCGGCGGCACACTGGGTCATCCCTGGGGCAATGCGGCGGGCGCAACTGCCAATAGAGTGGCACTGGAAGCGTGCGTTAAGGCACGTAACGAAGGACGCGCACTTATTGCAGACGGTCCCCAAATCTTACGTGAAGCAGCCGCACATTCCCCAGAATTAGCGGTCGCGATGGAAACATGGAAAGACATACGCTTCGAACAGCCAACTGTGGACACCCTGGACGCAACGCATGCGGTCTAA

Form IF-2 SSU:

ATGCCGTCGGAAACTTTTAGTTATCTTCCAAGCTTGACAGACGCTGAGGTACGGGCTCAGGCAGCAAACTTAGCCCGCGAAGGTCTGATCCCAGGCCTCGAGTATACACGTGAGCCGTCACCCCGGGACAGCTTCTGGAGCCTCTGGAAATTACCCTTCGTTGATCGTCCAAGTCCTGATGCCGTACTGGCGGAAATAGATCAGTGCGCAGCTGAGCATCCGGACGCTCATATA AAATTAGTGGGATATGACAGCGTGCGGCAGGGCCAGGTAGTGTCATTTGTGGTGCGCGTCCCACGT

Form IF-3 LSU:

ATGCAAAACGGACCTAACCACAAGGGACTGTTTGATGCCGGCGTGCACGAGTATCGCGAGAGTTACTATGACCCGGGTTATGTTCCGAAAGATACCGATTTTTTAGCCGCTTTTCGTGTGACGCCACAGCATGATGTCCCTCCTGAAGAAGCGGGTGCCGCTGTTGCCGCCGAGTCATCCACAGGGACATGGACCACGGTTTGGAGCGATCTTCTGACTGACATTGAACGTTATCGTGCGCGATGCTATCACATTGAACCCGTCCCTGGACAGGAGTCGCAGTTCCTGGCCTATGTTGCTTATCCACTGGATCTTTTTGAGGAGGGTAGCATCGTGAACGTGATGAGCTCCATTGTCGGTAACGTTTTTGGCTTTAAGGCTCTGAAGGCCCTGCGCCTGGAGGACCTTAGAGTACCTGTTGCCTACTTAAAAACCTTTCAAGGACCTCCACACGGTATCCAAGTCGAACGTGACCTGTTAAATAAGTATGGTCGCCCATTTCTCGGTGGCACAATTAAACCGAAACTCGGACTGTCCCCTCGCAATTATGGTCGTGCATGTTATGAATGTCTCCGTGGTGGTTTGGACTTTACAAAAGATGATGAGAACATTAACTCGCAGCCATTTATGCGCTGGCGTGACCGTTATTTGTTCGTGACCGAAGCCGTGAAAAAAGCTGAAGCGGAGACCGGCGAGCGCAAAGGTCATTACTTAAACGTCACTGCTCCAACCATGGAGGACATTTATGAGCGAGCAGAAGTCGCTCGGGATATGGGATCCCCAATTATTATGGTCGATTACCTTACAGTCGGATTCGCAGCCCATACAAGCTTGGCGAAATGGTGTCGTAAACACGGTATGCTTCTTCATGTGCACCGTGCTATGCACGCAGTAATTGATCGTCAGAAGAACCATGGTATTCATTGGCGTGTACTGGCAAAATGGCTCAGAATGGCTGGTGGCGACCATCTGCACAACGGTACCGTCGTTGGCAAATTAGAATCTGATCGCGGCTCTACTCTGGCAATCAATGATCTGCTGCGTAATGACTTTGTACCAGCAGATAAATCACGCGGAATATACTTTGATCAGCCCTGGGCGAGCCTGCCGGCGGTCCTGCCAGTAGCGTCCGGCGGTATTCATGTGTGGCACATCCCTGAATTACTTCACATATTTGGGGACGACGTTGTTCTGCAATTTGGAGGTGGCACCCAGGGTCATCCTGCAGGCAATGTGGCCGGTGCGACGGCCAATAGAGTGGCGCTGGAGGCGGCAGTGTTAGCGCGCAATGAAGGTCGAGATCTTGTCGCAGAGTCACAGGCTATATTGGTAGCTGCTGCCCGCCATTCACCCGAATTACGTGCTGCGTTAGATCTGTGGAAAGACATCGCCTTTGGTTTTGAGACGGTTGATACGCTGGATGCAGTGGTGGCGAAAGCTTAA

Form IF-3 SSU:

ATGAAACTGGAGACCTTTAGCTACCTCCCTCCTCTGACTCCACAACAGAAAGTGCGTCAAATTCAGTATATTCTGAACCAGGGCTTGATTCCCGGCGTTGAATATACCCGTCGTCCTGAGCCCCGTGATCATTACTGGCACATGTGGAAACTGCCGCTGTTCTCAGCACGCACTCCTGCGGATGTCATTCAGGAAGTCGAGGCATGCAAGGCAGAGAATCCTGGCACGTATATTA AACTGACCGGCTACGATAATAAACGGCAATGTCAGGTGATTTCCTTTGTAGTATACCAGCCCGATGAAAGA

Form IF-4 LSU:

ATGTCGGACTCTAATGAAAACGGCAAGGCACTGTTTGATGCGGGCGTGAAAGATTATTCAGCTGCGGGACGGTATTATTATGATATCGGGTTCAAACCAACAGAAACACAAGTTTTAGCTGCATTTCGAATCACGCCGCAGGAGGGCGTATCATTCGAAGAAGCAGCGGCCGCAGTTGCGGCCGAGTCTAGCTTTTCCACCTGGACCACAGTGTGGAGTGATTACCTCGTCGATGCCGCTCGGTACAGCGCTCGGACCTATGAAATGAAACCTGTCCCTGGTCATAAAGATCAGATGATGGCCTATATAGCGTACCCGCTTGATTTATTCGAAGAGGGCTCTATGCCCAATTTGATGTCGTCGATCGTGGGAAACGTTTTTGGCTTCAAGCCCTTAAGAGCCTTACGGTTAGAAGATATTTATTTTCCGCCTGCATTACTTACTACCTTCCAGGGGCCACCACATGGCATCCAGGTGGAACGCGATAAACTCGATAAGTATGGCCGCCCCCTGCTGGGCGCAACGATGAAACCAAAACTGGGCCTGAGTGCGCGTAATTATGGACGTTTGGTTTATGAAGCGTTAAGAGGAGGCCTTGATTTCACTAAAGATGACGAGAATATTACCAGTCAACCGTTTATGCGGTGGCGTGATCGCTTTGAATTTGTGATGGCCGCGGTAAAACAGGCTGAAGCTGAAACTGGCGAGCGTAAAGGTCATTACTTGAATGTTACTGCTGGAGATATGGAAGAAATGTACAGACGAGCTGAGCTTGCCAAAGAACTTGGCAGTCGAATTATCATGGTTGATTATCTGGTGGCGGGTTTCACTGCGTTCGCCTCTTTATCCAAGTGGTGCCGGGCGAATGGTATGCTGCTTCACGCCCATCGGGCGATGCATTCGGTTATGGATCGTCAACAAAATCATGGTGTACATTGGCGGGTGCTCGCAAAATGGTGCCGGATCGTAGGTGCAGATCATTTGCACAATGGTACAGTCGTTGGGAAACTTGAAGGAGATCGCGCCAGTACGATTGGCATTAACGAAATGATGAGAGCAACCGATGTTGCCAAAAAAGAAACATGCGCGGGTCACCACTTCGATCAACCCTGGCTCAGTATGCCCGCAATGTTCCCAGTCGCATCTGGTGGTATTCATGTTTGGCACATCCCTGAATTGGTTTCAATTTTCGGTGACGACGCAATCCTTCAATTTGGAGGGGGCACCGTTGGACATCCATGGGGTAGCGCCGCAGGAGCCACTGCGAACCGCGTGGCTCTGGAGGCCGTCATCCAGGCGAGAAACGAAGGAAGAGATTTACATACCGAAGGCACCGATATTCTGAAATCTGCTGCTAAGCACTCGGGAGAGTTGCGTGCCGCAATGGAAACATGGAAAGACATTACGTTTGAATATGAGAGTGTAGATGCCTTAGAAAAATCTTAA

Form I F-4 SSU:

ATGAAGACTGAAACCTTCTCCTATTTGCCGACGTTTACGCGTGAACAGGTGGAGAAACAAATACAATATTTCCTCAACAATTCTTGGGTTGTGGGGATTGAGTATACCAGTCAGCCGAATCCAAGTCTTGTATTCTGGGATTGGTGGCGTCTGCCGCTGTTTAACATGAAATCTGTTGGTGAAATTATGGCAGAGGTAGACGCATGTAAGGCGGCGAATCCTGATTGTTATATCCGT ATTACGAGCTATGATCATGTACGGCAAAGCCAGGTTATGGGGTTTGTGGTGCACCGAGT A

Form IF-5 LSU:

ATGGGTGAAACGAATGGGCAGGTTAGTCGTTTGTTCGATGCTGGAGTTAAAGACTATTCGGACGCGACGAGATACTATTACGATCCGGATTACGTCCCCGCCGAAACGGATGTGCTGTGTGCTTTTCGGGTGACCCCACAGCCAGGTGTTAGCTTCAAAGAGGCGGCTGCCAGCGTCGCTGCTGAATCATCAAGCGGAACCTGGACAACCGTTTGGACCGATCATCTGACCGATTTAAGTCGGTATAGTGCGAGATGCTACCGCATCGAACCAGTGCCGGGGCGTGATGACCAGTTCATTGCGTACATTGCCTATCCGATGGATCTTTTTGAAGAAGGTTCTGTTGTTAACATGATGTCGTCCATTGTCGGTAATGTATTTGGATTTAAAGCTATTCGTGCCCTGCGCTTGGAGGACGTAAGAGTTCCGGTTGCCTACCTTAAAACGTTTCAGGGTCCTCCCCATGGTATAGTAGTGGAAAGAGATCGCTTGAATAAATATGGAAGACCACTTCTGGGAGCCACTATTAAACCAAAACTGGGCCTTTCTGCAAAAAATTACGGCAGAGCGGTGTACGAGTCCTTGCGCGGTGGGTTAGATTTTACCAAGGATGACGAGAATGTGAATAGCCAGCCGTTTATGCGCTGGCGTGATCGATTTTTGTTTGTTATGGAAGCCGTACACAAAGCAGAATCAGAAACTGGAGAGCGTAAAGGACATTATCTCAATGTCACCGCTCCAACATTTGAACAGATGATCGAGAGAGCTGAAACCGCTCGTGAATTGGGCTCACGCATTATTATGGTTGATTTTTTGACAGCCGGATTTGCCGCTCACACGTCCCTGTCGCACTGGTGTCGTAAACATGGTGTATTGCTCCATTGCCACCGCGCTATGCACGCCGTCATTGATAGACAGCGCGATCACGGTATTCATTGGAGAGTTCTGGCCAAATGGGCCCGCATGGCAGGGTGTGATCACCTGCACAATGGCACTGTGGTAGGTAAGTTGGAAGGTGATCGGCAGTCAACGCTTGGCGTCAACGACCTGTTGCGTCTGGACGATGTGCCACAAGATCGTTCAAGAGGTATTTATTTTGATCAGCCGTGGGCTAGTCTGGCACCAGTTATGCCTGTGGCGAGTGGTGGTATTCATGTCTGGCATATGCCAGAGTTAGTACACCTTTTTGGTGATGACGCTGTACTGCAGTTTGGTGGTGGTACCCTTGGTCATCCTTGGGGCTCAGCTGCTGGAGCAACAGCCAATCGGGTGGCCTTAGAGGCAGTAGTACAGGCTCGCAATGCCGGACGGGACTTATTAGAGGAAGGTCCGGCGATACTGAAGCAGGCAGCACGAAGATCCCCAGAACTTCAACTGGCGCTGGAAACTTGGCAACAGGTGCGGTTTGACTACGACGTTGTTGACCGTTTAGATCCTGTCCATAGTGGTACCGAAGGATAA

Form IF-5 SSU:

ATGGCTCTTCGTACAGAGATGTTTTCATATTTACCACCAATGCCGCCTGAAGAAGTTCGTCAACAAGTAGAGTATCTTATACGCCGTGGTTATGTGCCCGGTATTGAGTTTACACAGCGCCTGGACAGCCATGACGACTTTTGGTCTTTCTGGAAATTACCGTTCTTTCGTGGGGCTACCGTGGACGGTGTTCTGGCTGAGCTGGAGGCGTGTAAAACGGCCCATCCAGGGGCAA CAATTCGCCTGACTGGCTATGATGCGCGACGGCAATGCCAGGTGCTGAGCTTCGTTGT GCACCGGCCGGCT

